# Predicting the feasibility of targeting a conserved region on the S2 domain of the SARS-CoV-2 spike protein

**DOI:** 10.1101/2023.03.05.531227

**Authors:** Pranav Garg, Shawn C. C. Hsueh, Steven S. Plotkin

**Affiliations:** Department of Physics and Astronomy, University of British Columbia, Vancouver, British Columbia V6T 1Z1, Canada; Genome Sciences and Technology Program, University of British Columbia, Vancouver, British Columbia V6T 1Z1, Canada

## Abstract

The efficacy of vaccines against the SARS-CoV-2 virus significantly declines with the emergence of mutant strains, prompting investigation into the feasibility of targeting highly conserved but often cryptic regions on the S2 domain of spike protein. Using tools from molecular dynamics, we find that this conserved S2 epitope located in the central helices below the receptor binding domains is unlikely to be exposed by dynamic fluctuations without any external facilitating factors, in spite of previous computational evidence suggesting transient exposure of this region. Furthermore, glycans inhibit opening dynamics, and thus stabilize spike in addition to immunologically shielding the protein surface, again in contrast to previous computational findings. Though the S2 epitope region examined here is central to large scale conformational changes during viral entry, free energy landscape analysis obtained using the path coordinate formalism reveals no inherent “loaded spring” effect, suggesting that a vaccine immunogen would tend to present the epitope in a pre-fusion-like conformation and may be effective in neutralization. These findings contribute to a deeper understanding of the dynamic origins of the function of the spike protein, as well as further characterizing the feasibility of the S2 epitope as a therapeutic target.

## Introduction

The rapid global spread of COVID-19 has provoked vigorous interest in studying the mechanisms of infection of the SARS-CoV-2 virus in order to develop therapeutic strategies. Attachment of the virus and its entry into the host cells is mediated by the trimeric spike (S) glycoprotein present on the viral surface.^1^ Spike is a prominent target of the adaptive immune responses raised during infection,^2,3^ and this feature, along with its essential role in ACE2-mediated cell entry^1,4–8^ has made it a primary choice for an immunogen in most COVID-19 vaccines used today.^9,10^

The spike protein undergoes several large-scale conformational changes as part of its function, and therefore its dynamics are as important to understand as its structure. Some of these dynamics are well-characterized: The receptor binding domain (RBD) has an up-down motion, wherein residues in the receptor binding motif (RBM) that bind ACE2 are only fully exposed in the up state.^11^ This conformational fluctuation-dependent exposure likely provides some degree of protection of epitopes from the immune system when the region is sequestered in the down state; nevertheless, the RBM is immunodominant.^3^ Following attachment, the S1 domain comprising the RBD and the N-terminal domain (NTD) separates from the S2 domain. The furin cleavage site between S1 and S2 is cleaved during viral biogenesis, ^1^ whereas cleavage at a second S2’ site just prior to entry activates fusion with the host cell membrane.^11^ There are two pathways of cleavage and subsequent fusion employed— an endosomal route involving cleavage at the S2’ site by cathepsin L, wherein the virus is first endocytosed before merging with the membrane of the endosome, and a direct entry route wherein the S2’ site is cleaved by TMPRSS2 and the virus merges at the cell membrane.^11^ The availability of two routes of cleavage and entry enables the virus to infect more cell types.^12,13^

After separation of S1 from S2 and cleavage at the S2’ site, the S2 domain undergoes a dramatic conformational change that allows it to perform fusion between the viral and host membranes. ^14^ The detailed mechanism and timing of the separation of S1 from S2 and the consequent conformational changes of S2 are not well understood, and may also differ between the endosomal and direct routes due to environmental factors such as the pH. The structure of the S2 domain following fusion has been determined, ^14^ but the short-lived intermediates where the S2 domain spans the viral and host membranes are not structurally characterized. The kinetic and thermodynamic properties of these changes are also currently not known.

Some insight into dynamics and conformational rearrangements of the spike protein have been obtained from cryo-EM and hydrogen-deuterium exchange (HDX) studies. Benton et. al.^15^ obtained structures of spike protein with multiple RBD’s either in the up state or bound to ACE2. They further observed that with multiple RBD’s bound to ACE2, the region of S2 at the top of the central helices, comprising residues 980-990, is exposed to solvent and has room to initiate extension of helix comprising residues 990–1035. This was proposed as the starting point for the formation of the membrane-spanning intermediate and embedding of the fusion peptide in the host membrane. ^15^

The same region of S2, comprising residues 978-1001 was highlighted in an investigation through hydrogen-deuterium exchange by Costello et. al.^16^ They found that the protein population has a bimodal mass distribution upon deuteration where state “B” exposes residues 978-1001 to the solvent to a greater degree than the classical prefusion state “A”. The interconversion between states A and B is slow, and which state is preferred (more stable) is temperature-dependant. Further, they found that the antibody 3A3 characterised by Huang et. al.^17^ binds an epitope within residues 980-1006. The antibody 3A3 was in fact found to be neutralizing in cellular fusion and pseudovirus assays.^17^

In a separate study characterizing a large library of nanobodies,^18^ one nanobody (S2-10) was determined to bind in the neighbourhood of residue 982 in the S2 domain, with synergistic effects when combined with an RBD-binding nanobody. That is, binding of RBD epitopes exposed in the up-state improved binding by S2-10. The authors also postulated that due to the smaller size, a nanobody may be able to access and bind this cryptic epitope more effectively than an IgG antibody.

The region of S2 encompassing residues 980-1006 is relatively well-conserved among the betacoronavirus lineage.^17^ The existence of conformational states where it is accessible suggests a promising target for the development of vaccines and antibodies that are robust against evolution and antigenic escape of the virus. Moreover, this site is central to large scale functionally important conformational changes—in the prefusion state, the residues 983 and 984 form the turn in a helix-turn-helix motif, which reconfigures to one continuous long alpha helix in the post-fusion state. A conformational state where this region is exposed may lie along the pathway to separation of S1 from S2 after protease-cleavage. Both the antibody 3A3^16^ and the nanobody S2-10^18^ are believed to achieve neutralization by blocking the region’s conformational transition to the membrane spanning fusion intermediate.

A common strategy to stabilize and increase the yield of class I fusion proteins in their prefusion conformation is to introduce prolines at the site of helix turn-extension transition.^19^ This strategy is beneficial for experimental studies as well as for vaccination, and has been extensively utilized for SARS-CoV-2. The “spike-2P”^20^ variant incorporates the mutations K986P and V987P, and the “hexapro”^21^ variant further includes proline mutations to F817, A892, A899 and A942. Importantly, the residues 986 and 987 that are mutated to proline are contained within this conserved sequence, and their mutation would be expected to affect antibody efficacy for this target. Indeed, the 3A3 antibody, which was raised against the spike-2P variant incompletely blocks infection by the native sequence^17^ and shows essentially no efficacy against the Omicron BA.1 strain.^22^ As well, the rate of interconversion between states A and B identified by Costello et. al. differs between the spike-2P and the hexapro variants,^16^ which suggests that the relevant dynamics that allow exposure of this S2 region are sensitive to mutations that stabilize the prefusion state.

Further, many of the vaccines deployed against COVID-19 use the spike-2P sequence,^10^ meaning that immune responses generated to the vaccine in this highly conserved region are actually tailored to an artificial sequence. These mutations can also have non-local effects on the binding of antibodies to other epitopes. An analysis of six neutralizing antibodies targeting the fusion peptide 816–843 found that the antibodies had reduced affinity for the spike 2P variant, and even lower affinity for the hexapro variant.^23^ Structural analysis of the bound structures showed that conformational flexibility was required for accommodating the antibody, which may explain why the stabilized mutants exhibited reduced binding despite the epitope being distant from the mutated residues.

While experimental analysis of the spike protein bearing a wild-type sequence in this region of interest is difficult, computational simulations may still allow us to gain insights.

Even if the “stem” and transmembrane regions of the spike protein are excluded from a simulation however, the system poses a considerable challenge to simulate due to its large size. The Folding@home project has utilized exascale computing to collect over 1 ms of adaptive sampling data^24^ for the reference sequence^25^ of the spike protein. Their simulations sampled states where the RBD is opened beyond what has been captured in the up-state of the cryo-EM structure (PDB 6VSB)^26^ (see Fig. S1A). While the probability density is low at these higher opening angles, the existence of the antibody CR3022^27,28^ that binds an epitope which is only exposed in a highly open state of the RBD suggests that other epitopes that are exposed in this open state, such as the neighbouring S2 region mentioned above, may also be viable targets.

Another 10 μs long simulation (10897850) from the D. E. Shaw group,^29^ starting with one RBD in the up state, opened up even further during the simulation, again exposing the above region of interest in the S2 domain (Fig. S1B). This simulation was performed using an initial structure that incompletely modeled the native protein however, in that several loops within the protein were missing, several glycans were incomplete, and as well, the sequence used in the simulations was the spike-2P sequence rather than the native spike sequence. A later simulation that corrected these shortcomings (11021571) did not sample such an open state. We note that these unbiased simulations are likely too short to deduce that the open state is present or absent in a significant fraction of spike molecules, but they do hint at the possibility of large-scale motions that would expose such a region.

An interesting question to consider in these dynamic processes is the role of glycans in allowing or disallowing motions that expose cryptic conserved epitopes. Casalino et. al.^30^ and Sztain et. al.^31^ have shown that dynamic motions of glycans not only conceal epitopes, but also modulate the relative populations of functionally relevant conformations. As well, Zimmerman et. al.^24^ found that the presence of glycans slightly biases the RBD towards open conformations, where the S2 region of interest is accessible. However, in these latter simulations, the force field used for the glycosylated system, CHARMM36, differed from that used for the unglycosylated system, AMBER03. Given the subtle differences in the probability distributions obtained for the states, a force field inconsistency has the potential to make interpreting this result problematic.

A goal of the present study is to examine the ease of accessing an open conformation of spike wherein the region of S2 that contains residues 980–990 is exposed, starting from a conformation where one of the three RBDs is in an up state. We present the potential of the mean force (PMF) along a simple angular reaction coordinate as the RBD is opened. We calculated this PMF for both fully glycosylated and unglycosylated variants of the spike protein, with an unmodified reference sequence,^25^ in order to understand the role of glycans in regulating this opening motion that exposes conserved parts of the S2 domain.

The region exposed in this open state is not only a conserved target for therapeutics, but also the site of extensive conformational change during membrane fusion. The experimental evidence summarized above indicates that it may indeed be accessible to therapeutic or serological antibodies. One way to exploit this property would be to immunize using a small fragment of S2 containing the relevant region, to direct immune response specifically to this conserved target. Since this region can exist in at least two different conformations, a designed vaccine antigen should account for their relative stability.

Therefore, we also calculated the free energy change profile for a conformational change between the pre-fusion helix-turn-helix conformation, and the single extended helix conformation that exists in the post-fusion state of the S2 domain, which is also expected to exist in the membrane spanning intermediate. In order to elicit antibodies that can neutralize viral entry by preventing a conformational change to the extended helix and thereby blocking membrane fusion, the putative vaccine would have the desired feature of exclusively presenting the pre-fusion conformation to the immune system. This might require designing scaffolds around the antigen in order to stabilize the pre-fusion conformation, without compromising the sequence identity of exposed residues.

To calculate the free energy cost of the conformational change corresponding to conversion of the helix-turn-helix to the straight helix, we used umbrella sampling along a reaction coordinate that connects the end states, but disallowed large deviations from either state. An approach described by Papoian and collaborators^32,33^ allows us to implement such a reaction coordinate for calculating the PMF. The method does not rely on knowledge of the true path between the end states, and incorporates restraints that prevent exploration of the vast phase space of unfolding conformations. The resulting PMF provides hints for antigen design and more generally the fusion mechanism of class I fusion proteins.

## Methods

### Simulation Systems

We calculated the free energy landscape of two different systems (Fig. 1 A,B). To study the ease of accessibility of the S2 region comprising residues 980–990, we simulated the “head” region of the spike protein, as follows. We started with the full-length spike protein structure in the RBD-up conformation, published by Casalino et. al.^30^ We trimmed this structure to only include residues 13–1146, removed glycans, and renumbered the chains to break the amide bond between residues 685 and 686 flanking the furin cleavage site. This starting structure was then imported into CHARMM-GUI^34^ and two structures were constructed, one lacking glycans, and one with glycans modelled according to tables S1-S3 from Casalino et. al.^30^ Disulfide linkages were made following the above-published structure. All chain termini, including at the furin cleavage site, were modeled in the corresponding charged states (NH_3_^+^ or COO^-^).

**Figure 1:**
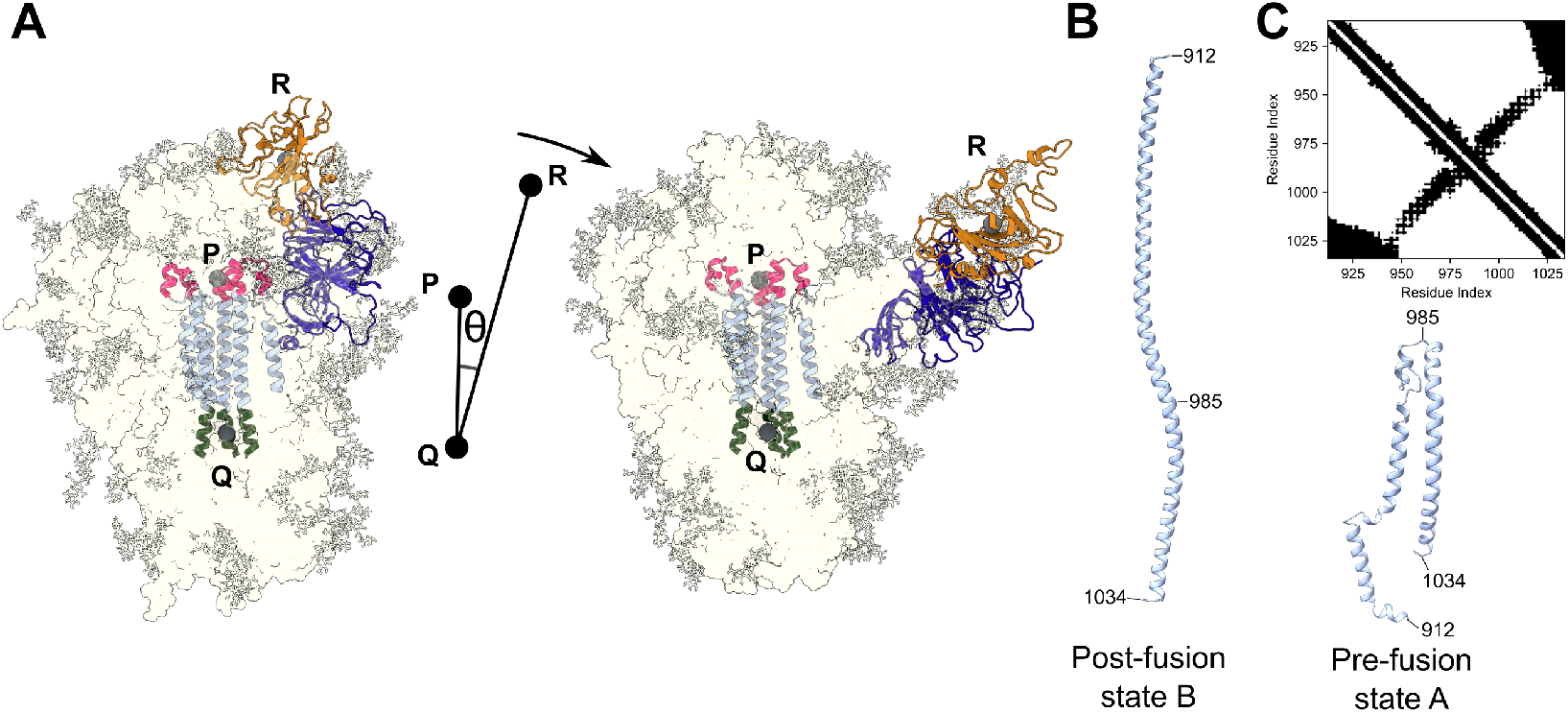
Simulation systems. (A) The head portion of the spike protein with one RBD in the up state. A portion of the S2 central helix region, the erected RBD, and its nearest NTD are shown using ribbons. Glycans are shown in licorice. The reaction coordinate is the angle *θ* determined by the CoM’s P,Q, and R (marked with gray spheres) of residue groups given in the main text. The NTD that moves with the RBD is shown in dark blue. An open state structure where the system has moved further along the reaction coordinate is also shown on the right. (B) Residues 912-1034 used in S2 helix extension simulation. These reference structures were obtained after equilibration of the pre-fusion bent state (right) and the post-fusion extended state (left). We assumed that the extended membrane spanning intermediate is in the post-fusion conformation. (C) A contact map of residue pairs used to define the reaction coordinate between pre- and post-fusion states in (B) (Eq. (1)). The matrix elements corresponding to the pairs included in the calculation are shown in black.

The calculation of the free energy change between the pre-fusion and extended helix states of the S2 core was performed with a monomeric system comprising residues 912-1034 (Fig. 1B). The extended helix conformation was taken from chain A of the post-fusion structure, PDB 7E9T.^14^ The pre-fusion state was modelled by extracting the relevant residues from chain A of the full-length spike protein structure in the RBD down conformation, published by Casalino et. al.^30^ Structures of both end states were capped with acetate and methyl groups on the N and C termini respectively, to remove end charge effects. Identical covalent bonding topologies were created for both states, containing the same number of water molecules and ions in order to allow interconversion between states generated from the two starting structures.

All simulations were performed in GROMACS-20 1 9.2^35^ patched with PLUMED 2.5.2,^36^ using the CHARMM36 force field.^37^ The proteins were placed in a dodecahedral box with periodic boundaries and solvated in TIP3P water^38^ with 150 mM NaCl. Solvation and ionization resulted in 1,649,564 and 1,575,556 atoms respectively for glycosylated and unglycosylated variants of the spike head system, and 578,529 atoms for the monomeric central helix system. During equilibration and umbrella sampling, the temperature was set to 310K, to emulate the active environment of the virus. Protonation states in the S2-extension system were consistent with an extracellular pH of 7.4, verified using the H++ server.^39^

Figures and numerical analyses used the NumPy, Pandas and Matplotlib packages for Python.

### Simulation scheme for S2 epitope accessibility simulations

To study the opening motion of the RBD beyond the up state, each of the two spike systems, one containing glycans and the other lacking them, was first equilibrated with position restraints on all heavy atoms. This comprised a 0.1 ns NVT simulation followed by a 0.2 ns NPT simulation, both with a 1 fs time step. Then we switched to a 2 fs time step, and conducted another 0.4 ns NPT simulation. Finally, the heavy atom restraints were removed, and the system was allowed to relax via MD in an NPT thermostat for 4 ns.

We were interested in studying the free energy barriers restricting the separation of the up-state RBD from the helix-rich region of S2 containing the heptad repeat 1 and the central helix. To do this, we calculated the potential of the mean force (PMF) along an angular coordinate *θ* = *∠PQR*, with coordinates *P, Q*, and *R* defined using the centers of mass (CoMs) of the C_α_ atoms of the following three groups of atoms: Residues 977-993 at the top of the central helices (P), residues 1020-1030 at the base of the central helices (Q), and residues 334-524 of the RBD (R) (see Fig. 1A).

Sampling of structures along this reaction coordinate was performed using replica exchange molecular dynamics coupled with umbrella sampling (REMD-US).^40,41^ Unlike temperature REMD,^42–44^ here all replicas are at the same temperature but have different restraints along a reaction coordinate, and exchanges are attempted between umbrellas that are adjacent along the reaction coordinate.

To obtain starting structures for the replicas involving RBD opening, we conducted a series of 2 ns umbrella sampling simulations starting from structures obtained after MD relaxation, which had a reaction coordinate angle of 0.26 rad (14.7°) and 0.28 rad (16.0°) respectively for the unglycosylated and glycosylated systems. In the direction of increasing θ, the bias center of the umbrella sampling started from 0.28 rad, and was increased by 0.03rad≈1.7° at the end of each simulation to start the next simulation in the sequential series, until a final angle of 1.09 rad (62.5°) was reached. In the direction of decreasing angle starting from the initial state, the bias center of the umbrella sampling started from 0.25 rad, and was decreased by 0.03 rad from one umbrella to the next, until it reached 0.10rad (5.7°). The spring constant for the harmonic restraint was 25,000kJ/mol/rad^2^. This series of simulations in both directions resulted in 34 umbrellas for each system.

In order to sample conformations similar to those explored by the D. E. Shaw trajectory 10897850,^29^ and the Folding@home trajectories,^24^ it was necessary to also restrain the NTD closest to the up-state RBD, so that the opening involves the combined motion of this NTD along with the RBD. This was implemented as an additional root mean squared deviation (RMSD) restraint in the 30 serial umbrella sampling simulations, with bias centers of the angle spanning 0.22–1.09 rad. The value of RMSD used in the restraint was calculated for the discontiguous group of atoms comprising residues 324–589 (residues 334–524 are shown in orange in Fig. 1A) of the up state RBD, and residues 13–305 of the neighbouring NTD (dark blue in Fig. 1A). This RMSD was calculated with respect to the native conformation at the end of the equilibration/relaxation trajectories of the glycosylated and unglycosylated systems. The RMSD was restrained with a spring of stiffness constant 10,000 kJ/mol/nm^2^ and a bias center of 0nm. The effect of applying this RMSD restraint is to preserve the relative position of the RBD and NTD to that in the native structure, while allowing those portions of the spike to collectively open, as in Fig. 1A. This RMSD restraint was not carried over to the REMD-US simulations used in the analysis; it was only applied during the biasing simulations to produce the initial conditions.

The REMD-US simulations were conducted on 18 NVIDIA Tesla V100 GPUs and 108 Intel Xeon Silver 4216 CPU over 14 days. The simulation time for each of the 34 replicas was 27.4 ns for the glycosylated system and 29.4 ns for the unglycosylated system. Systems in adjacent umbrellas were considered for exchange every 1 ps. Combined with the equilibration and initial configuration preparation, the cumulative simulation time was 1004 ns and 1072 ns for the glycosylated and unglycosylated systems respectively.

The potential of mean force (PMF) along the opening angle was calculated using the Multistate Bennett Acceptance Ratio (MBAR) algorithm, implemented in the PyMBAR package (v3.0.1).^45^ Convergence was assessed by dividing the reaction coordinate trajectory of each umbrella into four parts and using each of these parts to separately calculate the PMF (Fig. S2), and then checking if the PMF changed by more than one standard deviation over more than half the parameter space. Our simulations of the unglycosylated system did not reach convergence by the above criteria, nevertheless, the deviations are within 7k_B_T and do not affect the conclusion of glycosylation stabilizing the closed state. The final PMFs of both systems were calculated using the second half of the REMD-US sampling trajectories.

We also constructed PMFs for glycosylated and unglycosylated systems without the introduction of the additional RMSD restraint during preparation of the initial configurations (Section S1). This separate set of simulations sampled an alternative region of conformation space, where the NTD closest to the opening RBD remains close to the S2 helical core.

### Antibody accessible surface area of the S2 epitope

To better interpret the PMF obtained above, we wanted to assess the extent to which the conserved S2 epitope region is accessible to antibodies at various positions along the angular reaction coordinate. This antibody accessible surface area (AASA) was calculated for residues 737–765 and 961–1010, which is an extended neighbourhood of residues previously identified to be prone to exposure^15,16^ (see Fig. S1). We used a probe radius of 1nm, which is comparable to the complementarity-determining region (CDR) of an antibody chain (see e.g. Fig. 5 of ref.,^46^ and ref.^47^). The same portions of the trajectory that were used for the PMF calculation were also used for AASA analyses (i.e. the last 50% of REMD-US trajectories). A series of protein structures were extracted from each of the REMD-US replica simulations with an interval of 100 ps. The angle *θ* and the AASA were recorded for these frames, and the frames were binned by angle. The distribution of AASA values in each bin was then used to make box-plots showing the first and third quartiles, as well as the median and outliers. The AASA calculation was done using the MDTraj^48^ implementation of the Shrake and Rupley method.^49^

We also calculated the number of residues in the S2 epitope regions 737–765 and 961–1010 with non-zero exposure to antibodies for each frame in the last 50% of the sampled structures. Residues with AASA <1 Å (as probed by the antibody) were considered to have zero antibody exposure. A threshold of 0.1 Å has been used for this purpose in a previous study.^50^ In the structure of the antibody 87G7 bound to its RBD epitope (PDB 7N4l),^51^ the lowest per-residue AASA among the non-glycine residues Y421, L455, F456, F486 and Y489 is 3.6 Å, comparable to our 1 Å threshold. The frames were binned by angle in increments of 2.75°, and for each bin, the largest number of exposed residues is reported.

### Simulation Scheme for S2 Helix Extension

Systems at the two end states of the S2 helix shown in Fig. 1B were equilibrated using NVT and two NPT simulations, following the same recipe as the spike head systems (Section “Simulation scheme for S2 epitope accessibility simulations”). The last frames of the NPT simulations were used as reference structures for the end states and used to define the reaction coordinate. The method is not sensitive to the exact choice of the reference end states since all states in a basin around the end states are used for calculation of the free energy difference. Following Zhuravlev et. al,^32^ the similarity of two arbitrary protein conformations *X* and *Y* was defined as:

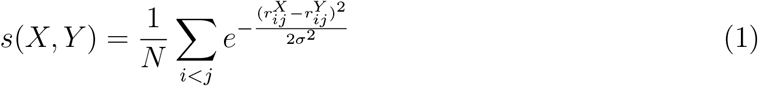

where *σ* =0.5nm and 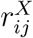 denotes the distance between atoms *i* and *j* in conformation *X*. The summation was restricted to those pairs of Cα atoms that were on non-consecutive amino acids, and either (i) were less than 1.4nm apart in either of the end states or (ii) were displaced by more than 10nm between the two end states. The former criterion preserves local structure and has previously been used with the same simulation technique.^33^ The latter inclusion criterion ensures that these largest motions directly contribute to the reaction coordinate calculation, and constrain structural exploration to a smaller portion of the phase space. See Fig. 1C for pairs of Cα atoms that satisfy these criteria. With the above criteria, the calculation of *s*(*X*, *Y*) depended on distances between *N* = 1784 pairs of atoms (Eq. (1)). The similarity *s*(*X*, *Y*) ∈ (0, 1], with *s*(*X*,*X*) = 1 for identical conformations, and lower values for dissimilar conformations.

Let *A* and *B* denote respectively the reference conformations for the extended helix, and the bent helix-turn-helix states (both shown in Fig. 1B). The position of a state *X* along the reaction coordinate ξ between these two states may then be defined as:

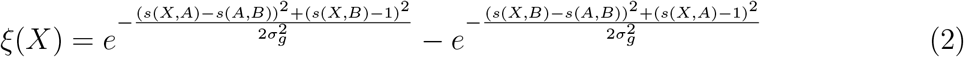

where *σ_g_* = 0.23. The similarity between end states *s*(*A*,*B*) was 0.486, and the reference states had reaction coordinates of ξ*_A_* = −0.993, ξ*_B_* = 0.993. (See Fig. 4).

**Figure 2:**
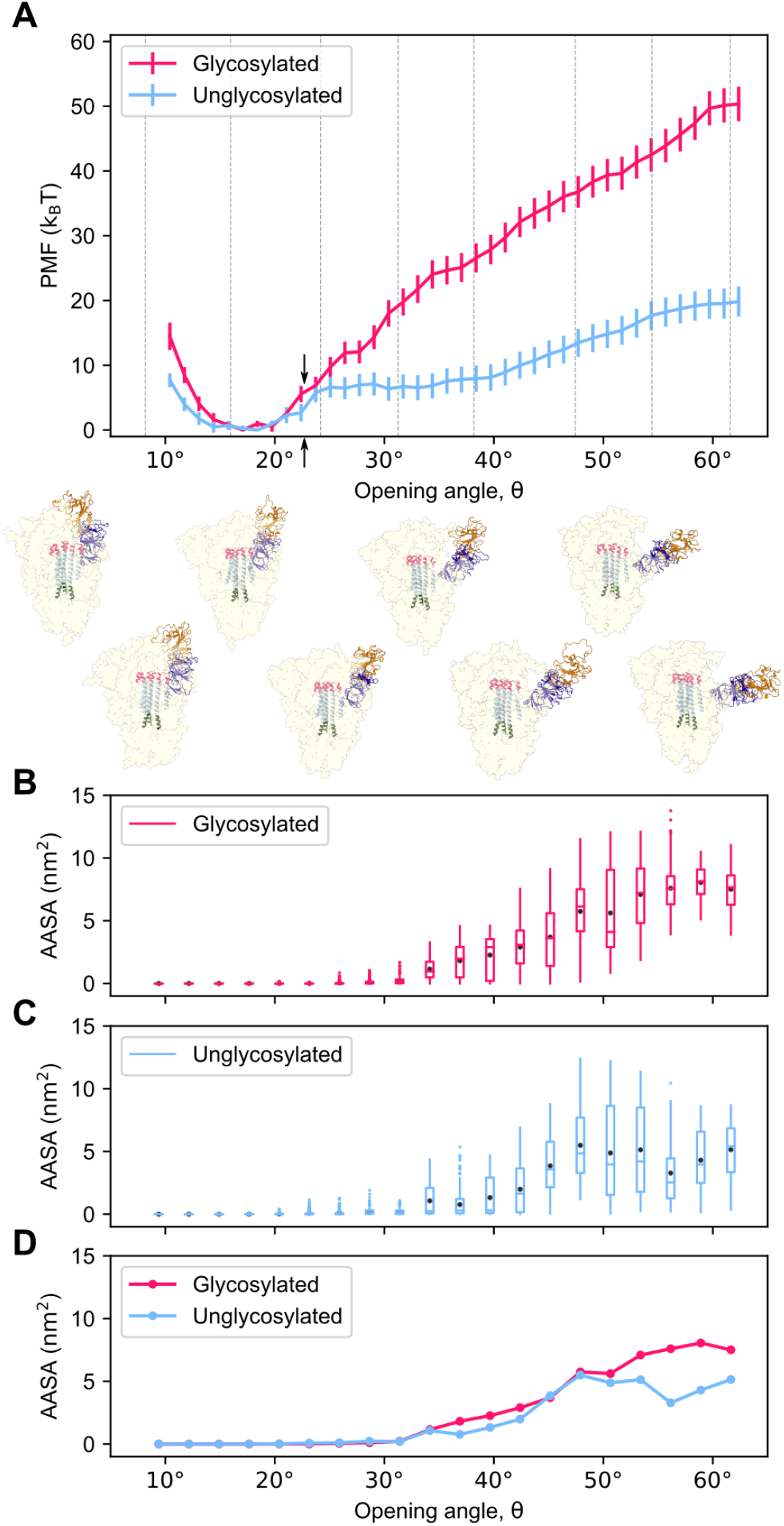
(A) The PMF of the glycosylated (dark pink) and unglycosylated (light blue) spike along the opening angle. Arbitrarily selected frames from the sampling are shown below, with corresponding angles marked by gray lines. The angle threshold of 22.7° is marked with arrows. (B,C) Box plots showing the AASA of the S2 region for glycosylated (dark pink) and unglycosylated spike (light blue) along the same coordinate as in panel A. Box boundaries correspond to the first and third quartiles, whiskers extend up to the last data-point within 1.5 times the inter-quartile distance, outliers are denoted as points, the medians as horizontal lines and the means as black dots. (D) The means from panels B and C are plotted together for comparison of the two systems. A plot of PMF against AASA is shown in Fig. S13A.

**Figure 3:**
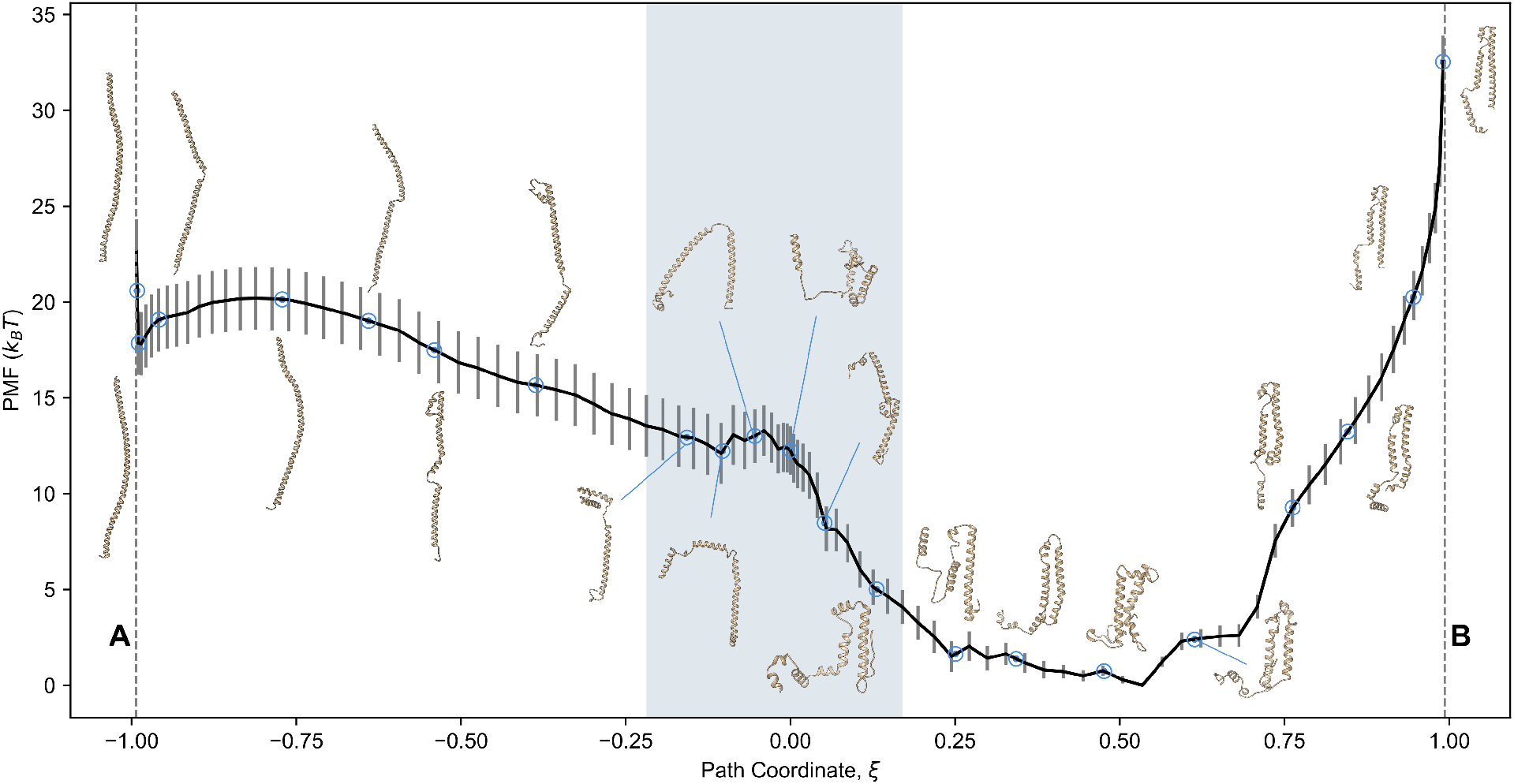
PMF calculated from S2 helix extension simulations. The extended structure is at ξ*_A_* = −0.993 and the pre-fusion bent “helix-turn-helix” structure is at ξ*_B_* = +0.993. Arbitrarily chosen structures from the umbrella sampling trajectories are annotated along the PMF to show how the structure changes along the reaction coordinate. The location of reference states A and B are shown as vertical dashed lines. The shaded region near ξ = 0 samples states with non-zero boundary potential (see Fig. S16).

**Figure 4:**
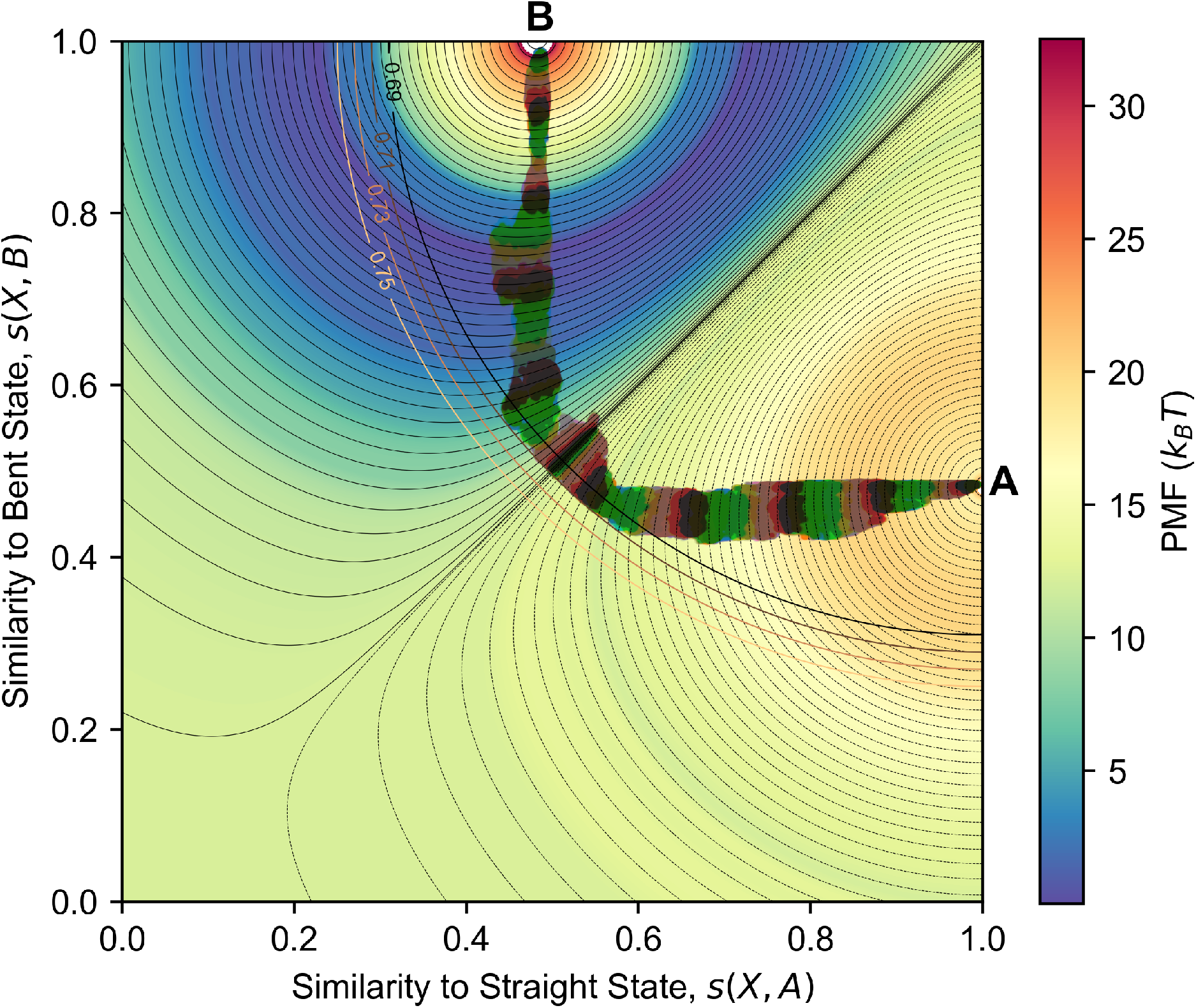
Phase space explored in helix extension simulations. The post fusion state A is located at *s*(*X*, *A*) = 1, *s*(*X*, *B*) = 0.486 and the pre fusion state B is at *s*(*X*, *A*) = 0.486, *s*(*X*, *B*) = 1. Umbrella centers along the coordinate ξ are shown as black lines. The circular arc with radius 0.69 centered at (1, 1) is the start of the repulsive radial boundary potential. Positions of structures sampled during the second half of the 16 ns umbrella sampling are shown as bands of various colours. The final PMF obtained (see Fig. 3) is shown in the background with a colour scale.

After equilibration, a moving restraint along this reaction coordinate was used to bias each end state to move towards the other. I.e. the extended helix structure, which had ξ = −0.993, was pulled using a moving umbrella that started at ξ = −0.993 and ended at ξ = +0.993. Similarly, the bent helix structure was pulled towards ξ = −0.993. These simulations also implemented a temperature annealing scheme. The temperature was first raised from 310K to 450K over 0.2 ns without moving the restraining umbrella. This temperature was maintained over the next 4 ns where the restraining umbrella was continuously moved towards the opposite state, before finally being cooled to 310K over 0.2 ns (see Fig. S3 A, B). Pulling at a higher temperature reduces the likelihood of the protein remaining trapped in local minima along the path, which would have hindered its ability to follow the moving restraint. In practice, the spring constant for the moving restraint was set to 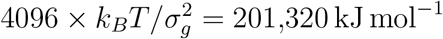; other sufficiently large values should also work as well.

During the pulls between end point conformations, coordinates of the system were recorded every 0.5 ps. A series of locations along the reaction coordinate were chosen for umbrella sampling (Fig. S3 C,D). These locations were spaced closely at the end states as well as in the region near ξ = 0. The latter was because the region ξ ≈ 0 encompasses a relatively vast region of conformational space, and we wanted to ensure adequate sampling in this region. Near the end points, we found that pulling simulations failed to reach close to the opposite end state than the one they started from, likely due to the steep energy landscape at those positions (c.f. Fig. 3) that would need increased sampling, or possibly indicating a shortcoming of the reaction coordinate definition in applying sufficient biasing forces near the end points. This non-uniform spacing of umbrella centres was implemented by first choosing 100 evenly spaced numbers *x_i_* between ξ(*A*) = −0.993 and ξ(*B*) = 0.993, and then transforming them to the reaction coordinate values ξ*_i_* using the function

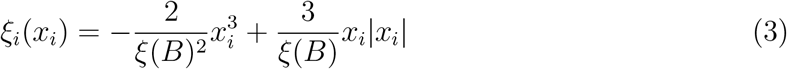

(plotted in Fig. S4). In addition, we included umbrellas at ξ*_i_* = ±1 to increase sampling at the end states. The spring constants for each umbrella was set to a value determined by its distance Δ*x* from the neighbouring umbrella towards ξ = 0. For a given umbrella with separation Δ*x*, the spring constant was set using

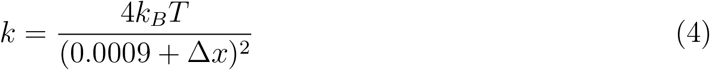

where *k_B_T* = 2.577 kJ mol^-1^, which gave spring constants in the range 1.1 × 10^4^ kJ mol^-1^ to 4.6 × 10^6^ kJ mol^-1^. Spring constants closer to the center of the transformation at ξ = 0 have stronger spring constants according to the Lorentzian function in Eq. (4). It is possible that with a more appropriate choice for the spread of the gaussian, *σ_g_* in equation Eq. (2), we could have avoided the use of unequally spaced umbrella centers. This choice of spacing and spring constant resulted in adequate overlap between umbrellas however, so it is not expected to affect the calculation of the free energy difference.

With the ξ*_i_* values in Eq. (3) chosen as spring centers, umbrella sampling simulations were initiated by extracting the conformations closest to these ξ*_i_* from the two pulling trajectories. The starting frames for the three umbrellas closest to each of the two end states (6 umbrellas total) were chosen exclusively from the pulling trajectory that started at the corresponding end state. The rest of the starting frames in the umbrellas were chosen by alternating between the two trajectories.

The resulting 102 umbrellas were then simulated in parallel replica exchange molecular dynamics umbrella sampling simulations (REMD-US),^40,41^ all at 310K, but with different umbrella potentials. Replica exchange was attempted every 1 ps. To begin, in addition to the restraining potential, all replicas were pulled for 1 ns along a direction defining the shortest distance between their location in *s*(*X*,*A*), *s*(*X*,*B*) space, and the straight line path that joins the end states (solid black line in Fig. S3 E). The force is applied along this direction using the shortest distance of each point to the line in Fig. S3 E, with a spring between those two corresponding points of spring constant 1.0 × 10^4^ kJ mol^-1^. Trajectories showing the decrease of the shortest distance to the line in Fig. S3 E indicate that increasing the 1ns time will not substantially increase the proximity to the straight line, indicating that physical constraints prevent the transformation trajectory from approaching the straight line between the two end state conformations.

After this initial pull, a boundary potential was implemented to prevent trajectories from exploring states near *s*(*X*,*A*) = 0 and *s*(*X*,*B*) = 0, far from the end states (i.e. deviating towards the bottom left in Fig. S3 E). Deviating slightly from the treatment in Zhuravlev et. al.,^32^ instead of a tangent hyperbolic wall we used the following quartic boundary potential, defined using the distance *d*(*X*) in (*s*(*X*,*A*), *s*(*X*,*B*)) space from the point *s*(*X*,*A*) = 1, *s*(*X*,*B*) = 1:

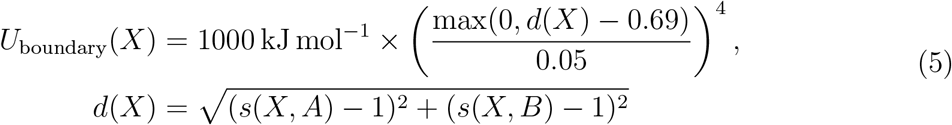

This potential is shown in Fig. S5.

By observing the distribution of frame positions in the (*s*(*X*,*A*), *s*(*X*,*B*)) phase space in the absence of this potential (Eq. (5)) when frames were pulled towards the straight line path (Fig. S3 F), the radius of the bounding potential in *s*(*X*,*A*), *s*(*X*,*B*) space was set to 0.69 in practice. At this radius, only 10.1 % of the ensemble eventually obtained crosses into the region of non-zero boundary potential. A smaller radius would potentially confine the trajectories too much and reduce the extent of interconversion between states.

With this “guard-rail” potential in place, the REMD-US umbrella sampling with 102 replicas was run for 18 ns in each umbrella, of which the first 2 ns was excluded from analyses. The total useful sampling time summed over all umbrellas was 1.63 μs. The position ξ of the trajectories along the reaction coordinate was recorded every 0.02 ps. Histograms of these positions along ξ were checked for sufficient overlap between adjacent umbrellas (Fig. S6). Using the time-correlation function of the positions ξ, the positions were down-sampled to an uncorrelated subset for accurate uncertainty estimation^45^ using scripts in the PyMBAR package (v3.0.5), and then used to construct the PMF along the reaction coordinate, *G*(ξ). The PMF profile of four equal 4 ns parts of the 16 ns trajectories was compared to judge convergence (Fig. S7). Informed by the convergence plots, the final PMF was calculated using the latter half of the trajectories, i.e the latter 8 ns of this ensemble.

## Results and Discussion

### Direct separation of the RBD from the central helices is energetically unfavourable, and glycans protect against this separation

Fig. 2A shows the PMFs calculated from REMD-US trajectories of the glycosylated and unglycosylated models of the spike protein. Energetic penalties for opening one RBD along the angle *θ* are steep for both glycosylated and unglycosylated variants. In the absence of glycans, only 0.74 % of the protein ensemble is predicted to obtain an opening angle of at least 22.7°, while in the presence of glycans, this fraction decreases to 0.077%. The angle 22.7° (marked with arrows in Fig. 2A) is a relevant threshold since it is the opening angle in the structure of an ACE2-bound RBD, PDB 7A95,^15^ and opening further would not be required for the attachment function.

The minimum free energy along the opening angle occurs at 17.1° for the glycosylated system and 18.4° for the unglycosylated system. For comparison, in the experimentally resolved structure 6VSB,^26^ the up-state RBD has an opening angle of 21.0°, while the relaxed and equilibrated structure of the glycosylated spike protein provided by Casalino et. al.^30^ that was used as an initial structure in our study has a lower opening angle of 14.7°. The flatness of our calculated PMF and thus low energetic cost of angular fluctuations in this neighbourhood of angle may explain this apparent disagreement between the experimentally determined and simulated structures. The range of angles for which our PMF is less than 3k_B_T is 13.6°–21.3° for the glycosylated system and 12.3°–22.5° for the unglycosylated system.

Due to the RMSD restraint used during generation of the initial configurations, the system begins exploring its allowable phase space starting from structures where the up state RBD and the NTD closest to it move together, reproducing conformations similar to those seen in the D. E. Shaw group trajectory 10897850,^29^ as well as trajectories from the Folding@home group (see Fig S2 of ref.^24^). An equilibrium ensemble generated over much longer simulation times might include states where the motion of the RBD and the NTD is not coupled. In order to sample such conformations, we conducted a separate set of simulations without an RMSD restraint (Section S1), and we obtained another PMF (Figs. S8, S9).

In this new set of simulations, the NTD remained attached to the helix-rich core of the spike protein while the RBD was pulled away. The energy penalties for opening the RBD beyond the up state in this ensemble are somewhat larger than those observed in simulations where the NTD moves along with the RBD, both with and without glycans. In this ensemble, the range of angles for which the PMF is less than 3k_B_T is 12.4°–22.2° for the glycosylated system and 13.2°–24.6° for the unglycosylated system, slightly wider than that in PMFs in Fig. 2A. While the relative contributions of these two motions (opening just the RBD, or both the RBD and the NTD) cannot be compared without much more exhaustive sampling, we note that neither pathway allows much unassisted opening at physiological temperature. Moreover, along both pathways, glycans increase the energetic penalties for this opening motion.

A potentially interesting role of the difference in PMF for glycosylated and unglycosylated spike could be to aid fusion through the endosomal pathway of entry. In the digestive environment of the endosome, glycans are degraded.^52^ The change to a flatter energy land scape with facilitated RBD separation may then allow the S1 domain to more easily detach from the S2 fusion machinery.

The range of residues included in our model, 13–1146 is comparable to the range 27–1147 used in both prior computational works mentioned above,^24,29^ but our model of the protein differs in other ways. Zimmerman et. al.^24^ used a different force field (AMBER03) for their unglycosylated system than for their glycosylated system, which used the CHARMM36 force field. Moreover, based on the structures provided on https://osf.io/fs2yv/, the glycosylated structure is not cleaved at the furin site. The authors observe a large range of motion in their trajectories (i.e. the opening angle *θ* ranged 7° to 46° for glycosylated spike and 9° to 48° for unglycosylated spike), however convergence tests were not performed. In the case of D. E. Shaw trajectory 10897580,^29^ the structure is missing 149 residues, typically those in disordered loops. Unlike our simulations and those of ref.,^24^ the Shaw trajectory sequence also contains the spike 2P mutations K986P and V987P, and glycans are modelled as single sugar molecules instead of full-length branched chains of sugars, and 6 of the 22 residues that are known to be glycosylated^53^ are not modelled as such.

The Folding@home group also obtained a distribution of the position of the RBD in glycosylated and unglycosylated systems (see Fig. S1 of ref.^24^), which can be quantitatively compared against our PMFs when projected on the same reaction coordinate (Fig. S10). Zimmerman et al. find that the probability distribution of RBD deviation from its down-state position was broader with glycans present, allowing more of the protein to adopt open conformations. In contrast however, we found here that the unglycosylated spike has a broader distribution of opening and S2 exposure (Fig. S10C). However, the RBD deviations observed in our simulations are generally larger than those observed by the Folding@home group, which indicates that the sampling in the two studies occurs in significantly different regions of the phase space. In particular, all of our simulations proceeded from an up-state conformation, and did not sample down-state conformations, while the Folding@home simulations started from a down-state conformation (PDB: 6VXX),^1^ and sampled both down-state and up-state conformations. Thus, we can argue that, given a fluctuation brings an RBD to an up-state, glycans will *hinder* the further opening of the RBD, while the Folding@home study argues that the presence of glycans *facilitates* the down-to up-state transition, and the consequent exposure of the S2 domain due to that transition.

### The cryptic S2 epitope around residues 980-990 is exposed in open states of the spike protein

To understand the implications of the calculated PMFs for immunogenicity, we calculated the antibody accessible surface area (AASA) for residues 737–765 and 961–1010 (“the S2 epitope”) along the opening angle, *θ* (see Section Accessible surface area analysis for S2 epitope accessibility simulations). Antibodies typically require binding to at least 3–5 residues^54–56^ to specifically recognize an epitope. We therefore identified the range of opening angles at which this requisite number of residues starts to be accessible.

For glycosylated spike, an up-state RBD needed to be biased to an angle of about 25° before the REMD-US simulations began to sample snapshots that had more than 1 nm^2^ (1nm^2^ = 100Å^2^) total AASA (outliers of the box plot around 25° in Fig. 2B). This corresponds to an energy cost of 9.6 k_B_T (Fig. 2A). It was also observed that at 30.0°, the number of S2 epitope residues with non-zero antibody exposure increased to 5 (Fig. S11). The corresponding free energy cost to access this angle in the absence of any additional facilitating factors was prohibitively high, about 16.9 k_B_T. I.e., the probability to open to an angle >30.0° is ≈1 × 10^-6^ %. In the absence of glycans, the S2 epitope was accessible at lower angles: the REMD-US simulations sampled snapshots with >1nm^2^ AASA at about 22^°^ (outliers of the box plot around 22^°^ in Fig. 2C), corresponding to an energy cost of 2.6k_B_T (Fig. 2A). It was observed that at 20.4°, the number of S2 epitope residues with non-zero antibody exposure increased to 5 (Fig. S11). This angle was accessible at a low free energy cost of 1.6k_B_T. I.e., the probability to open to an angle >20.4° is ≈6%.

An ACE2-bound RBD, as seen in PDB 7A95, has an opening angle of 22.7^°^, meaning that even this functionally unavoidable level of opening would have allowed the region to be targeted by S2-directed antibodies if the protein were unglycosylated. Further, the PMF shows that the energy penalty for opening an unglycosylated spike is much lower than when glycans are present, meaning that a greater proportion of cryptic epitopes would be susceptible to antibodies. Glycans therefore confer protection both through direct shielding and by modulating large scale dynamics, which can have protective effects beyond those involving the residues on the outer protein surface.

The glycans that confer protection from exposure of the S2 epitope are those linked to N61, N165, N234, T323, and N343 on the chain containing the moving NTD. Exclusion of all other glycans from the AASA calculation leaves the plot of AASA against opening angle essentially unchanged (Fig. S12). Glycans on N165 and N234 were also identified by Casalino et. al.^30^ for their role in stabilizing the RBD in the up state. They observed that these glycans insert themselves into the space underneath the RBD, reducing its flexibility. This insertion may also be responsible in protecting the S2 epitope which is directly underneath.

The AASA of the S2 epitope in the unglycosylated system shows a decreasing trend between 48° and 57° (Fig. 2C,D). Inspection of REMD-US trajectories in the umbrellas near this angle revealed that the motion of the RBD was not coupled with the motion of the NTD closest to it. Instead, the NTD drifted back to a position that was in close proximity to the S2 core, thereby shielding the S2 epitope. This contribution to epitope shielding can be seen in the protein image in Fig. S14B, where omission of NTD residues 13-305 and the associated glycans on this chain led to an increase in the ΔAASA between 48° and 57° (Fig. S14C). This effect was not observed in the glycosylated system (Fig. S14A), possibly because the bulky glycans hindered the interaction between the NTD and S2 core. One consequence of glycans inhibiting protection by the NTD was that the unglycosylated system had lower AASA than the glycosylated system for the S2 epitope (Fig. 2D). The presence of glycans has previously been observed to inhibit interaction between protein domains in other simulation studies. The glycan on N90 of ACE2 interferes with binding to the RBD,^57^ and the glycan on N234 on spike inhibits the transition of the RBD from the up state to the down state.^30^

In simulations initiated without constraining the neighbouring NTD to move with the up state RBD (Section S1), we found that for a glycosylated system, an up-state RBD needed to be pulled to about 45° before the S2 epitope started to be accessible to antibodies (Fig. S8B). The free energy cost for such a degree of opening was prohibitive, about 43 k_B_T. This indicates that for glycosylated spike, the pathway involving coupled motion of the NTD and RBD is more relevant for S2 epitope exposure, i.e that combined detachment of RBD and the nearest NTD from S2 core has a lower free energy cost to reveal the cryptic S2 epitope. For both glycosylated and unglycosylated system in these alternative simulations, the averaged AASA values along the opening angle are generally lower than those observed in the trajectory involving the combined motion of the RBD and the NTD (Fig. S8D). Further, in the unglycosylated system, the RBD is more permissive to opening than in the glycosylated system (Fig. S8A), similar to the trend observed along the pathway where the NTD moves along with the RBD.

Our calculation of the PMF was based on the native sequence rather than the spike-2P variant often used in experiments. We found that separation of the RBD from the central helices along an angular coordinate is highly unfavourable in its natural glycosylated state. While experimental studies have indicated that the S2 epitope underneath the RBD is accessible to antibodies, our finding casts doubt on an unassisted opening mechanism without the aid of any exogenous interactions, wherein the natural flexibility of this multi-domain protein would result in a vulnerable sub-population that can be neutralized by an antibody binding at the exposed site. Since structural experimental studies using the native spike sequence in purified full-length spike protein are difficult, one way to better test this result would be to immunize animals with a small construct containing the epitope of interest, and test the elicited antibodies in neutralization assays with pseudoviruses^58^ expressing the native spike protein.

Despite the relatively high degree of conservation in the S2 epitope in the broader context of betacoronavirus spike sequences, at the time of this writing, a few mutations in this region have arisen in lineages that have had widespread geographical prevalence. The B.1.1.7 lineage (alpha) had mutation S982A, and the lineage BA.1 (omicron) had mutation L981F, which are however no longer widely prevalent at the time of writing.^59^ Consistent with this, the S2-directed antibody 3A3 was measured to have lower IC_50_ against the alpha variant compared to the original Wuhan sequence.^17^ An ideal characterization of efficacy in targeting the S2 epitope would require checking the strength of antibody binding to a library of possible mutants in the S2 epitope. Even mutations outside the region could modulate susceptibility to antibodies via allosteric effects.

### Extension of a monomeric fragment of the S2 central helix is energetically unfavourable

Fig. 3 shows the PMF along the reaction coordinate ξ connecting the bent pre-fusion conformation of the selected S2 region and the extended conformation that is believed to be shared by the membrane spanning intermediate and the post-fusion state. Sampling of conformations in the two dimensional *s*(*X*, *A*), *s*(*X*, *B*) phase space is shown in Fig. 4, together with the PMF obtained. The expected form of such a PMF for a protein with two observed “end-point” conformations is two basins separated by a potential barrier. The observed form of the PMF for our system has a very narrow and shallow basin near the extended state at ξ ≈ −0.986, and a wide basin towards the bent state. The local minimum near the extended state has a free energy of 17.8 k_B_T higher than the global minimum, indicating that the extended state is not the most favoured conformation for this contiguous protein fragment. Note that the rapid increase in PMF near the end points is due to conformations constrained to have a high similarity to the reference states, which reduces their entropic contribution to the free energy. Near ξ ≈ 0 however, there are a relatively large number of conformations available to the protein.

The free energy basin defined by 0.04 < ξ < 0.78 has free energies below 10k_B_T. The structures in this basin have a Boltzmann-weighted similarity of 〈*s*(*X*, *B*)〉 = 0.73 to the reference bent state structure (at ξ*_B_* = +0.993) and 〈*s*(*X*, *A*)〉 = 0.47 to the reference extended state structure (at ξ*_A_* = −0.993). The Boltzmann-weighted root-mean-squared deviations of these states from the reference structures for the bent and extended states are 1.52nm and 4.64nm respectively (Fig. S15). I.e. while the system prefers a bent state overall, the most favoured conformations of the isolated peptide deviate considerably from the bent state structure observed in the context of the whole spike protein.

The PMF profile indicates that the system favours a state that is bent rather than an extended state, but the global minimum also shows deviations from the native prefusion conformation in the context of spike protein, as mentioned above. Using such a monomeric fragment as a vaccine antigen would therefore be predicted to present a prefusion-like conformation, but whether the antibodies raised would also bind the S2 epitope in its native context with significant affinity would need experimental validation.

A similar investigation for hemagglutinin revealed that the same transition from a bent to an extended state is energetically unfavourable for the influenza virus as well.^60^ There are differences in the system and methodology between our study and that in ref.^60^ however—in the previous study, the pH was low (4.0 vs. our value of 7.4 to mimic the extracellular environment) to emulate the late endosome; the protein fragment simulated was trimeric rather than monomeric as in our study; and the experimental pre-fusion state in hemagglutinin had less helical content than the prefusion state of SARS-CoV-2 spike. Lin et al.^60^ obtained a free energy difference of about 5–10 k_B_T between the prefusion and post-fusion conformations for hemagglutinin. That is, in both SARS-CoV-2 spike and influenza hemagglutinin, the extension of the central helix is not driven by an inherent instability in this portion of the protein, a feature that we speculate could be shared with other class I fusion proteins. We note that the simulated numbers could change significantly in the presence of membrane, which was not included in either the previous study of Lin et al. or our present study.

The technique used here to define the reaction coordinate ξ is adapted from Zhuravlev et. al.,^32^ and is indifferent to the system and conformational change involved. While the free energy difference calculated is independent of path, the structures sampled by the intermediate umbrellas may not be representative of the biologically realized pathway between the states. Further, due to the presence of the bounding potential (Eq. (5)), the PMF profile away from the end states is not indicative of the actual transition pathway. In particular, the locally increased free energy near ξ = 0 is likely artificial since the trajectory encroaches into the repulsive boundary in that region of the reaction coordinate (see Fig. S16 and Fig. 4). Following the rationale of previous studies,^32,33^ we do not pursue a reweighting scheme for the boundary potential because the free energy differences between states that are in the region of zero boundary potential (Fig. S16) are not affected by its presence. Attempting to correct the PMF profile accounting for the boundary potential would require sampling the prohibitively large phase space near ξ = 0.

## Conclusion

Targeting a conserved portion of the spike protein could potentially enable vaccines and therapeutics that are less susceptible to evasive mutations. Here, we investigated the feasibility of targeting an S2 region containing the predominantly conserved residues 980-990. This region is also targeted by the antibody 3A3 and the nanobody S2-10. We calculated the PMF along an angular coordinate for the process of separating the RBD from the spike core, which exposes this S2 region. This calculation was based on the native sequence rather than the spike-2P variant used in many structural studies. The calculation was performed for the isolated protein in solvent, so it effectively calculates the likelihood of a spontaneous opening. We found that separation of the RBD from the central helices is highly unfavourable in the spike protein’s natural glycosylated state, but is more feasible in the absence of glycans. The shielding of the cryptic S2 epitope by glycans is thus achieved both through changes to protein dynamics as well as through direct physical protection. This finding essentially rules out an unassisted mechanism leading to exposure of the S2 epitope, and adds to the ways in which glycans are important to survival of the virus.

In order to inform the development of a stable antigen for immunization containing this conserved target, we quantified the energy landscape of helix extension at this site, which leads to a membrane spanning conformation. We found that for a monomeric fragment, the helix prefers a bent conformation by approximately 18 k_B_T. If such a protein fragment were expressed and used as an immunogen, it should be expected to adopt the pre-fusion-like bent conformation for presentation to the immune system. This finding also means that the energy driving this conformational change is not derived from the inherent instability of each helix-turn-helix motif, but rather from other interactions (e.g. spike-membrane interactions) in the context of the full spike structure, a finding that may also be applicable more broadly to class I fusion proteins.

Beyond exploring the therapeutic potential of a conserved epitope, our study also provides insights on the functional mechanisms of the spike protein. Both reaction coordinates explored here, for RBD separation and helix extension, are relevant to the large-scale conformational changes that occur during membrane fusion.

## Author Contributions

*Conceptualization:* P.G. and S.S.P.; *Methodology:* P.G and S.C.C.H.; *Software:* P.G., S.C.C.H.; *Data analysis:* P.G., S.C.C.H. and S.S.P.; *Visualization:* P.G. and S.C.C.H.; *Resources:* S.S.P; *Writing—original draft:* P.G. and S.C.C.H.; *Writing—review and editing:* S.S.P.; *Supervision:* S.S.P.; *Funding acquisition:* S.S.P.

## Acknowledgement

This research was supported by the Digital Technology Supercluster, Canadian Institutes of Health Research Transitional Operating Grant 2682, Compute Canada Resources for Research Groups RRG 3071, and UBC ARC Sockeye Advanced Research Computing (https://doi.org/10.14288/SOCKEYE, 2019). P.G. and S.C.C.H. have received support from MITACS Accelerate Scholarships. We also thank Mark Thachuk for allowing access to the former Westgrid Orcinus computer cluster.

## Supporting Information Available

The following files are available free of charge.

- PDF: supplementary Methods and Figures

## TOC Graphic

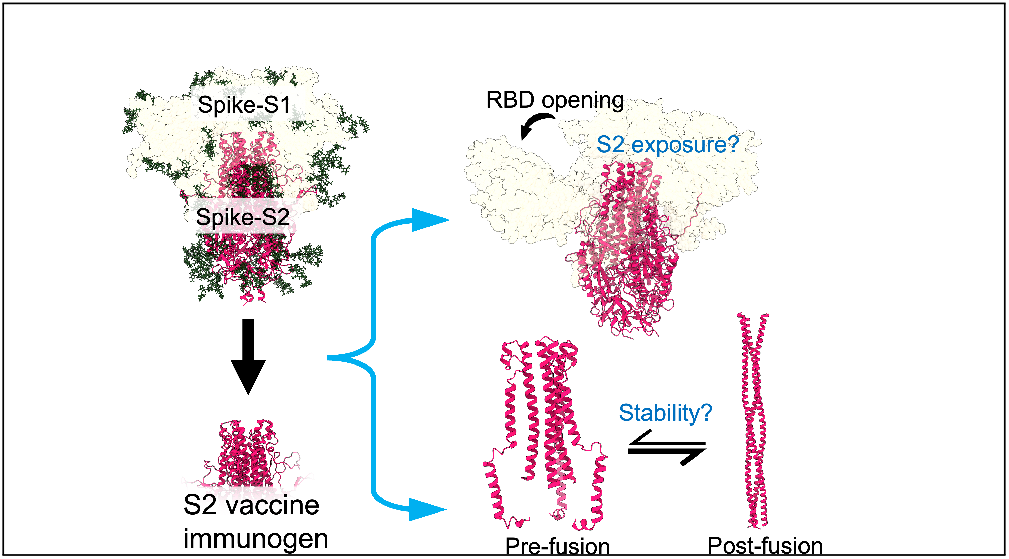

Some journals require a graphical entry for the Table of Contents. This should be laid out “print ready” so that the sizing of the text is correct. Inside the tocentry environment, the font used is Helvetica 8 pt, as required by *Journal of the American Chemical Society.*

The surrounding frame is 9 cm by 3.5 cm, which is the maximum permitted for *Journal of the American Chemical Society* graphical table of content entries. The box will not resize if the content is too big: instead it will overflow the edge of the box.

This box and the associated title will always be printed on a separate page at the end of the document.

