## supplementary material for "Predicting the feasibility of targeting a conserved region on the S2 domain of the SARS-CoV-2 spike protein"

### S1 Simulation scheme for S2 epitope accessibility simulations for uncoupled NTD/RBD motion

While our primary set of simulations for epitope accessibility involved constraining the NTD closest to the RBD to open along with the RBD during the initial biasing, we also calculated the PMF if no such constraint was introduced.

Starting from the equilibrated structure for the glycosylated and unglycosylated systems, a series of 34 short 2 ns umbrella sampling simulations were generated for each system in directions of increasing angle and decreasing angle, in order to obtain a starting structures for generating a PMF. In the direction of increasing angle, the position of the center of the umbrella started from 0.26 rad (14.7°) for the unglycosylated system and 0.28 rad (16.0°) for the glycosylated system, and was increased by 0.03 rad≈1.7° at the end of each 2 ns simulation to start the next simulation in the series, until an angle of 1.09 rad (62.5°) was reached for the glycosylated system and 1.10 rad (63.0°) was reached for the unglycosylated system. In the other direction, the position of the center of the umbrella started from 0.23 rad (13.2°) for the unglycosylated system and 0.25 rad (14.3°) for the glycosylated system, and was decreased by 0.03 rad at the end of each 2 ns simulation to start the next one in the series, until an angle of 0.10 rad (5.7°) was reached for the glycosylated system and 0.11 rad(6.3°) was reached for the unglycosylated system. The spring constant for biasing was 25 000 kJ/mol/rad$^2$. The 34 umbrellas were then sampled independently (instead of the parallelized REMD-US scheme described in the main text).

At various stages during the umbrella sampling, the potential of mean force (PMF) along the opening angle was calculated using the Multistate Bennett Acceptance Ratio (MBAR) algorithm, implemented in the PyMBAR package [1]. Convergence was assessed by dividing the reaction coordinate trajectory of each umbrella into four parts and calculating the PMF using the corresponding segments of all umbrellas. If the PMF profile calculated from the last two quarters of the trajectories deviated by more than an error bar for more than half of the PMF data points in angle space, the non-converged subset of umbrellas were sampled further. The choice of umbrella sampling simulations to extend was made using the Mann-Kendall trend test [2]. Only umbrellas with an increasing or decreasing trend in the position along the reaction coordinate were continued further. The calculation and simulation extension were repeated until the PMFs constructed by the last two non-overlapping portions of trajectories were mutually within the error bar of each other (Fig. S9).

At this stage, the cumulative simulation time across the 34 umbrellas was 768 ns for the glycosylated system, and 940 ns for the unglycosylated system. The longest duration of an individual umbrella were 35 ns and 68 ns respectively for the glycosylated and unglycosylated systems.

For the unglycosylated system, this produced a converged PMF profile in the last two quarters of the simulation time (Fig. S9 B). The final PMF was calculated using the last half of the sampling from each umbrella (Fig. S8 A). However, for the glycosylated system, this process did not yield a fully converged PMF (Fig. S9 A). So we implemented an umbrella sampling simulation coupled with replica exchange molecular dynamics (REMD-US), where systems in adjacent umbrellas were considered for exchange every 1 ps. This parallel sampling of all 34 umbrellas was conducted for 9.8 ns. The final total sampling time for the glycosylated system was 1.1 µs, comparable to that of the unglycosylated system. The reaction coordinate trajectory from the REMD sampling was divided into three parts to calculate the convergence of the PMF. The convergence was assessed to be adequate (Fig. S9A). The final PMF was then calculated using the last 2/3 of the REMD sampling.

The same portions of the trajectory that were used for PMF calculations were also used for antibody accessible surface area (AASA) analyses (i.e. the last 66 % of REMD-US for glycosylated spike and the last 50 % of ordinary umbrella sampling for unglycosylated spike).

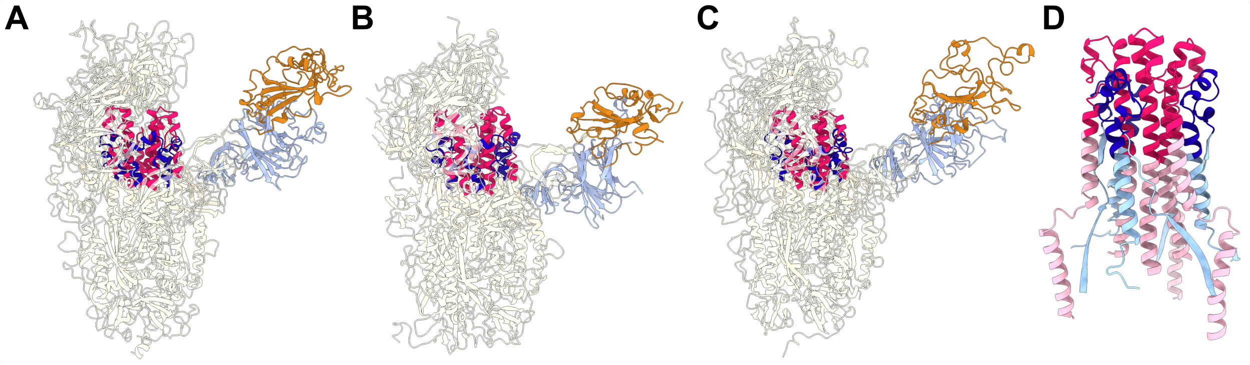

Figure S1: Region of interest on the S2 domain.
In all three panels, residues 961-1010 are shown in dark pink, and residues 737-765 are shown in dark blue. In panels A and B, the RBD residues 334-524 and NTD residues 16-305 are in orange and light blue respectively. (A) A frame from the trajectory of glycosylated spike published by the Folding@home group [3] (B) A frame from trajectory 10897850 published by D. E. Shaw group [4] (C) A frame from our glycosylated spike trajectory involving the opening of an RBD along with its closest NTD. (D) Closeup of residues 720-790 and 920-1033. Residues outside 961-1010 and 737-765 are shown in light pink and light blue.

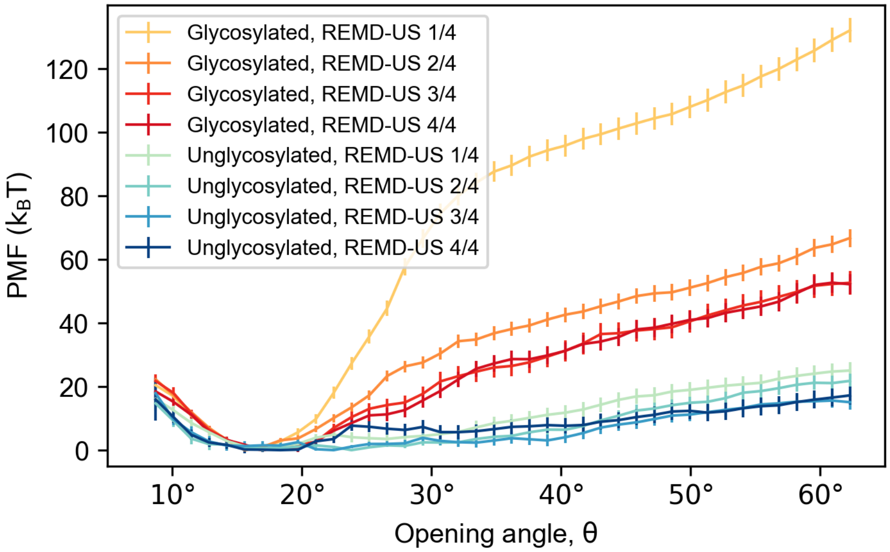

Figure S2: Convergence analysis for the S2-exposure REMD-US simulations.
Different non-overlapping portion of the umbrella sampling trajectories were used to construct PMFs. Fractions denote the portion used (e.g. 2/4 means second of four segments). (A) The Glycosylated system achieved convergence in the second half of the trajectories, where the PMFs are mutually within one standard deviation of each other. (B) The unglycosylated system shows small deviations in all four PMFs.

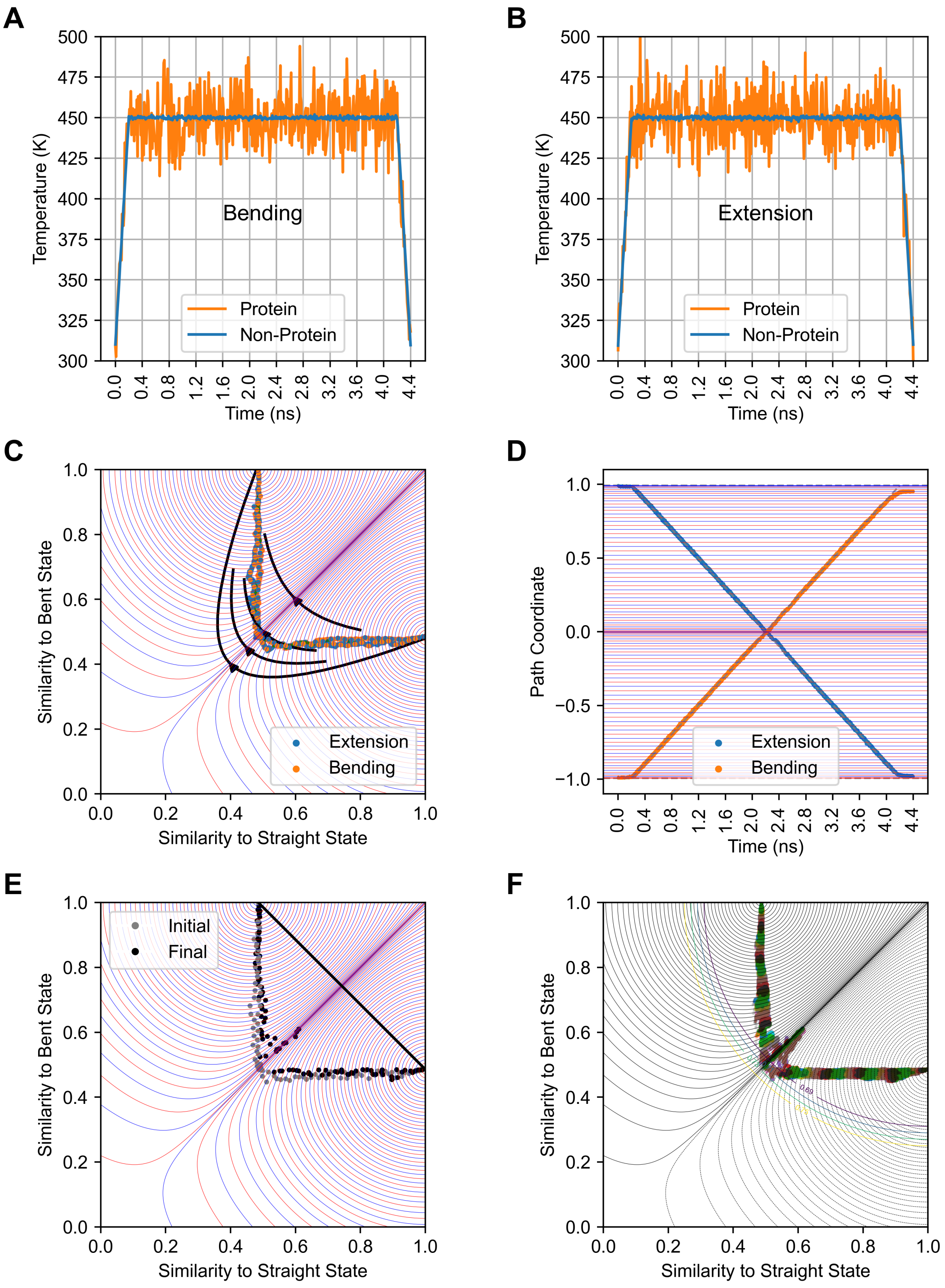

Figure S3: Generation of intermediate structures along the reaction coordinate for the S2-extension simulations.
(Caption continued overleaf)

Figure S3: (Continued from previous page.)

Panels A and B show the temperature profile for the pulling simulations starting at the bent and extended states respectively. The protein and non-protein (water molecules and ions) were coupled separately to the temperature reservoir.
Panels C and D show the trajectory of the pulling simulations starting from the bent (blue) and extended (orange) states. In panel C, the trajectory is plotted along the 2D phase space of similarity coordinates $s(X, A)$ and $s(X, B)$ and the progress along the time coordinate is excluded. Black lines indicate the approximate direction of steepest-descent paths of decreasing $\xi$, and are a guide to the eye. In both panels, blue and red lines indicate values of the coordinate $\xi$ chosen to be spring centers during umbrella sampling. Spacing of the centers as well as spring constants are non-uniform (see Methods in the main manuscript). Points plotted in panel C indicate starting structures closest to each chosen umbrella center, used to initiate further simulations. Starting structures selected from the pulling trajectories were subjected to a spring force towards the straight diagonal line shown in panel E. Positions of the frames before and after this 1 ns simulation are shown as grey and black points respectively. Panel F shows the distribution of structures sampled during this simulation. This distribution was used to set the radius for the boundary potential to be 0.69 (see equation 5 of Methods).

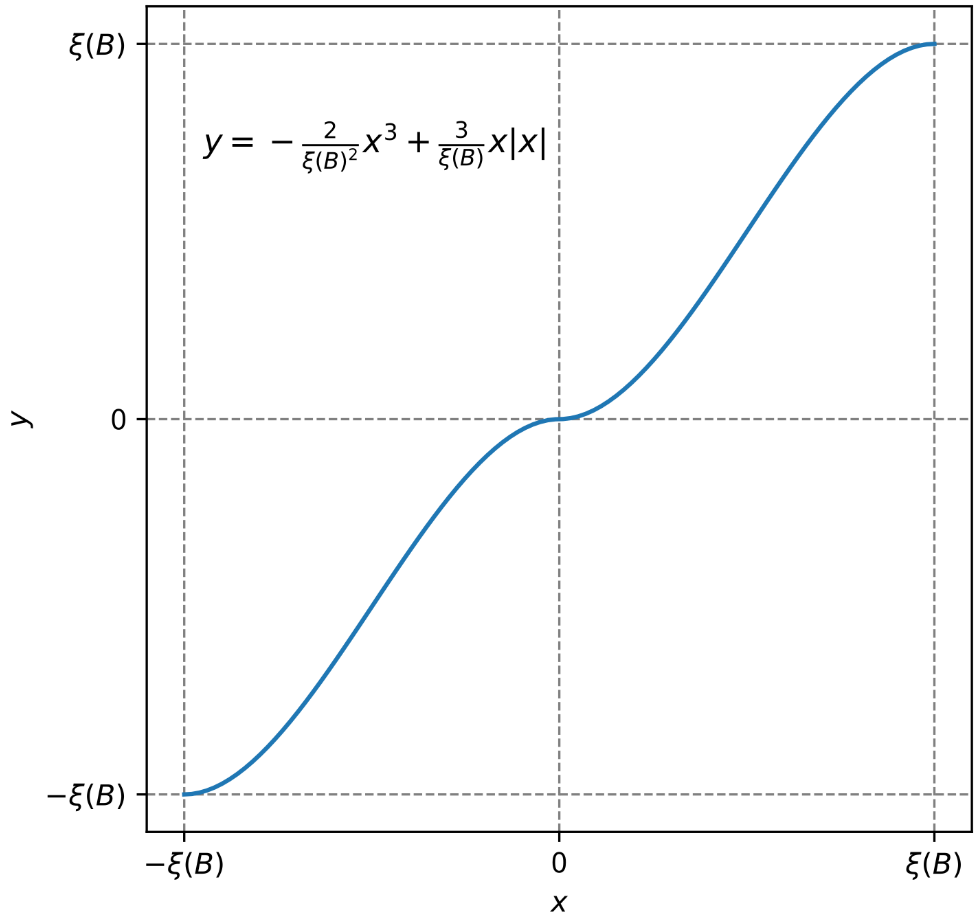

Figure S4: Spacing function for selecting umbrella sampling centers (Eq. (3) in the main text).
$y$ here indicates the positions of the umbrella sampling centers along the reaction coordinate. Uniformly spaced points along $x$ are transformed to points $y$ with in increased density near the end points and near $\xi = 0$.

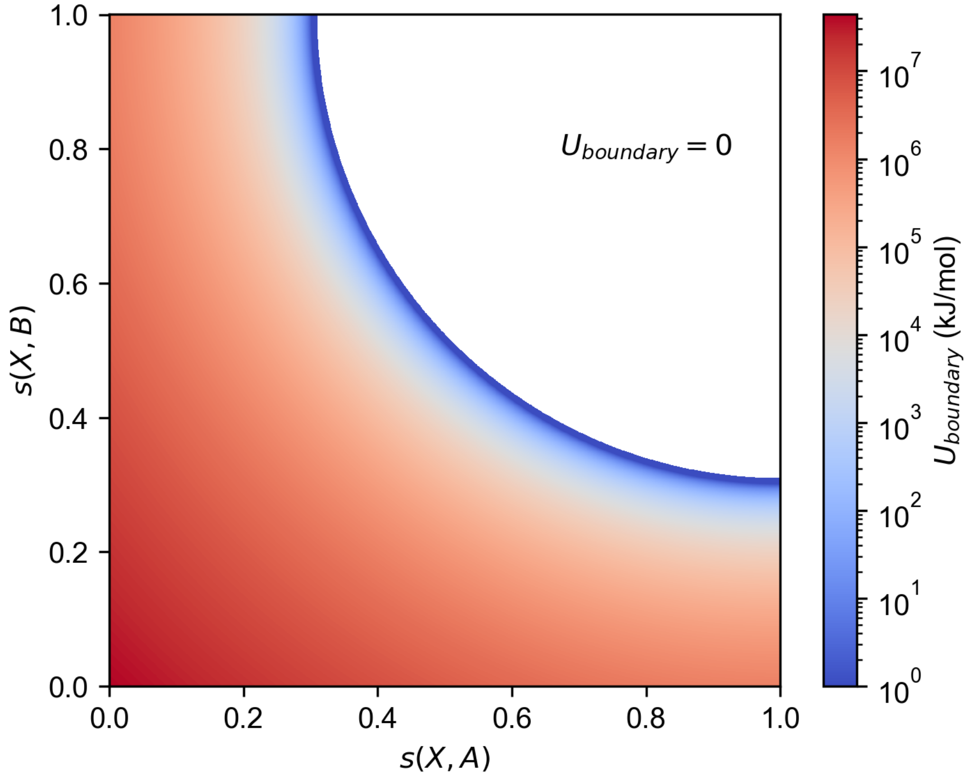

Figure S5: Boundary potential.
A plot of the quartic potential $U_{\text{boundary}}(X)$ defined by Eq (5) in the main text. Note that $k_B T \approx 2.6\,\text{kJ}\,\text{mol}^{-1}$.

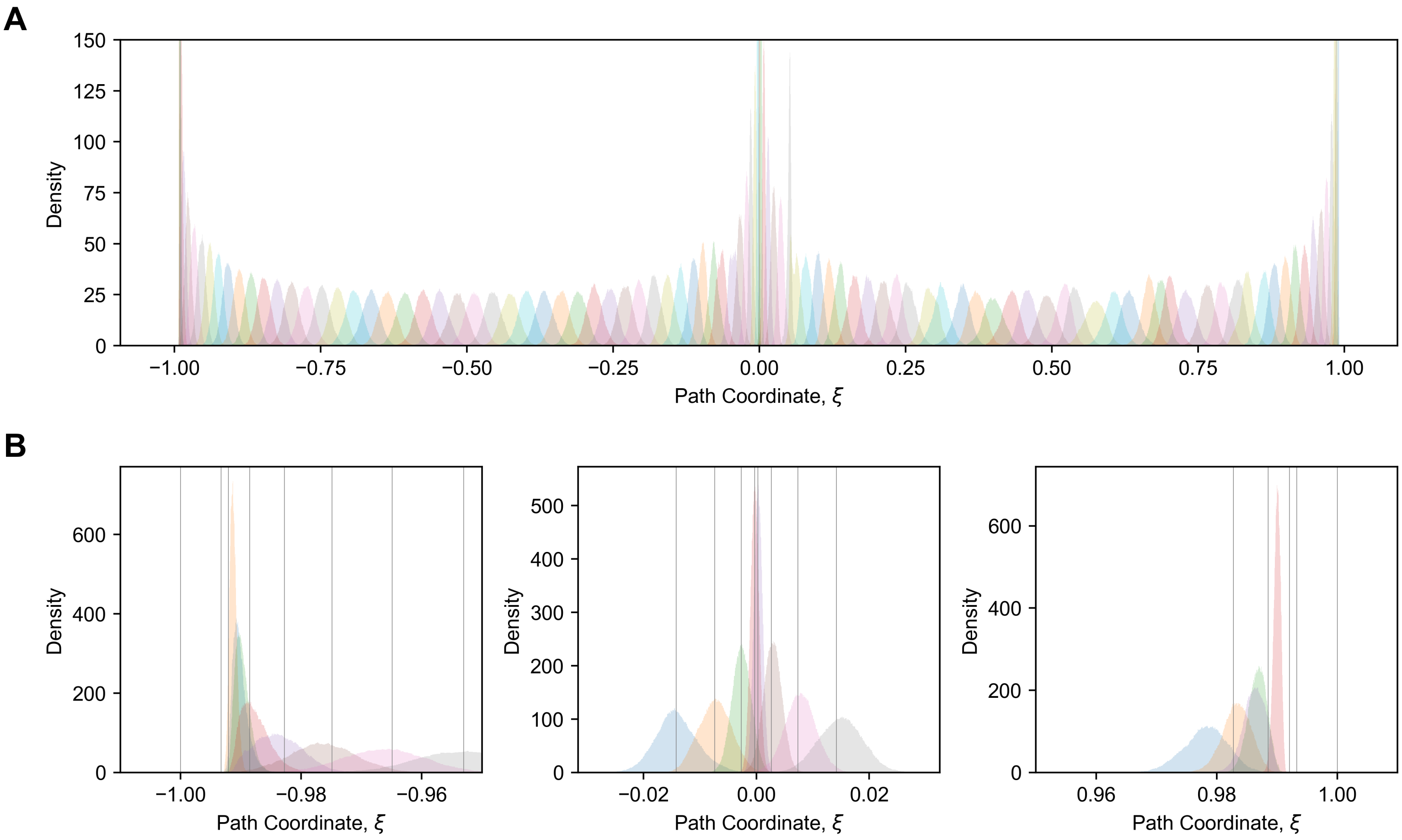

Figure S6: Umbrella sampling of structures during S2-extension simulations.
Positions of structures sampled during the 12 ns umbrella sampling simulations are visualized as histograms along the reaction coordinate $\xi$. Umbrellas were closely spaced near $\xi = 0$ and the end points $\xi = \pm 0.993$. The corresponding histograms are clipped on the y-axis in panel A and shown in detail in panel B. Vertical lines in panel B show the positions of umbrella centres. The large deviations of histograms from these centres is reflected in the steep increase in PMF near the end points.

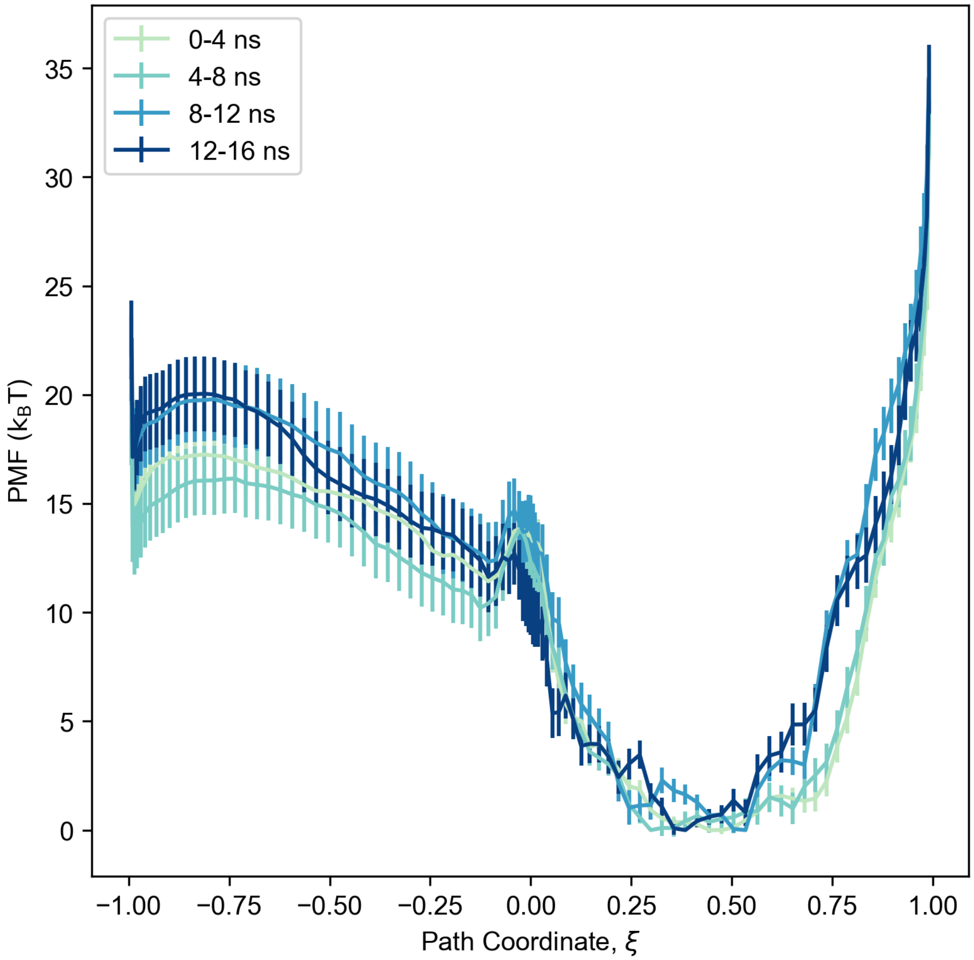

Figure S7: Convergence of the PMF in S2-extension simulations.
The 16 ns production trajectories were divided into four equal parts (see legend). Each part was individually subsampled to generate an uncorrelated subset of positions along the path coordinate, and then used to calculate the PMF.

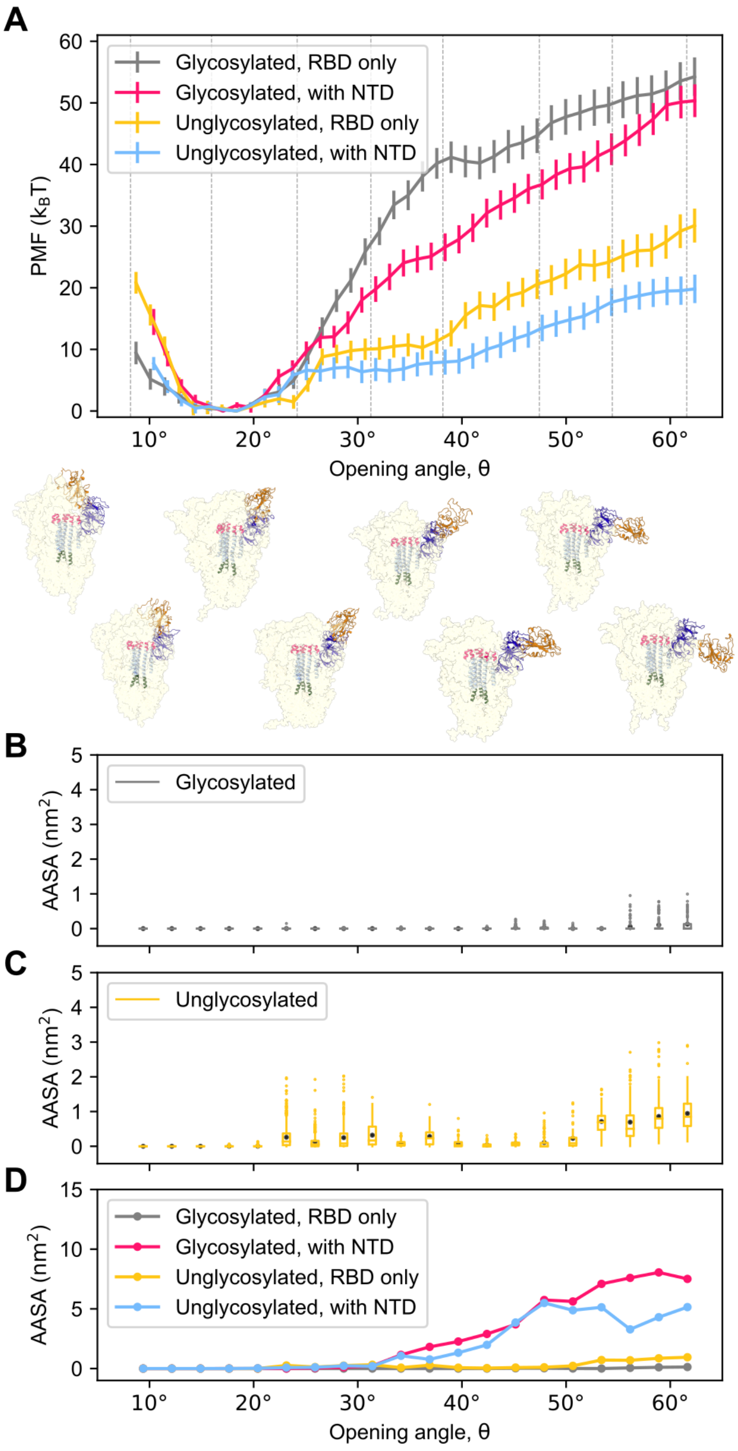

Figure S8: The PMF and AASA analysis for RBD opening simulations, for both NTD-coupled motion, and the alternative S2-exposure simulations where the RBD was pulled independently of its nearest NTD.
(Caption continued overleaf)

Figure S8: (Continued from previous page.)

This figure is analogous to Fig. 2 in the main text. (A) The PMF of the glycosylated (gray, pink) and unglycosylated (yellow, blue) spike along the opening angle. Arbitrarily selected frames from the sampling are shown below in order of increasing angle left to right, with corresponding angles marked by gray lines. The PMFs from the simulations involving combined motions of the RBD and NTD are shown in pink and blue for comparison ("with NTD"). (B,C) Box plots showing the AASA of the S2 region for glycosylated (gray) and unglycosylated spike (yellow) along the same coordinate as in panel A (RBD only). Box boundaries boxes correspond to the first and third quartiles, whiskers extend up to the last data-point within 1.5 times the inter-quartile distance, outliers are denoted as points, the medians as horizontal lines and the means as black dots. (D) The means from panels B and C are plotted together for comparison of the two systems, together with data from Fig. 2D of the main text. A plot of PMF against AASA is shown in Fig. S13B.

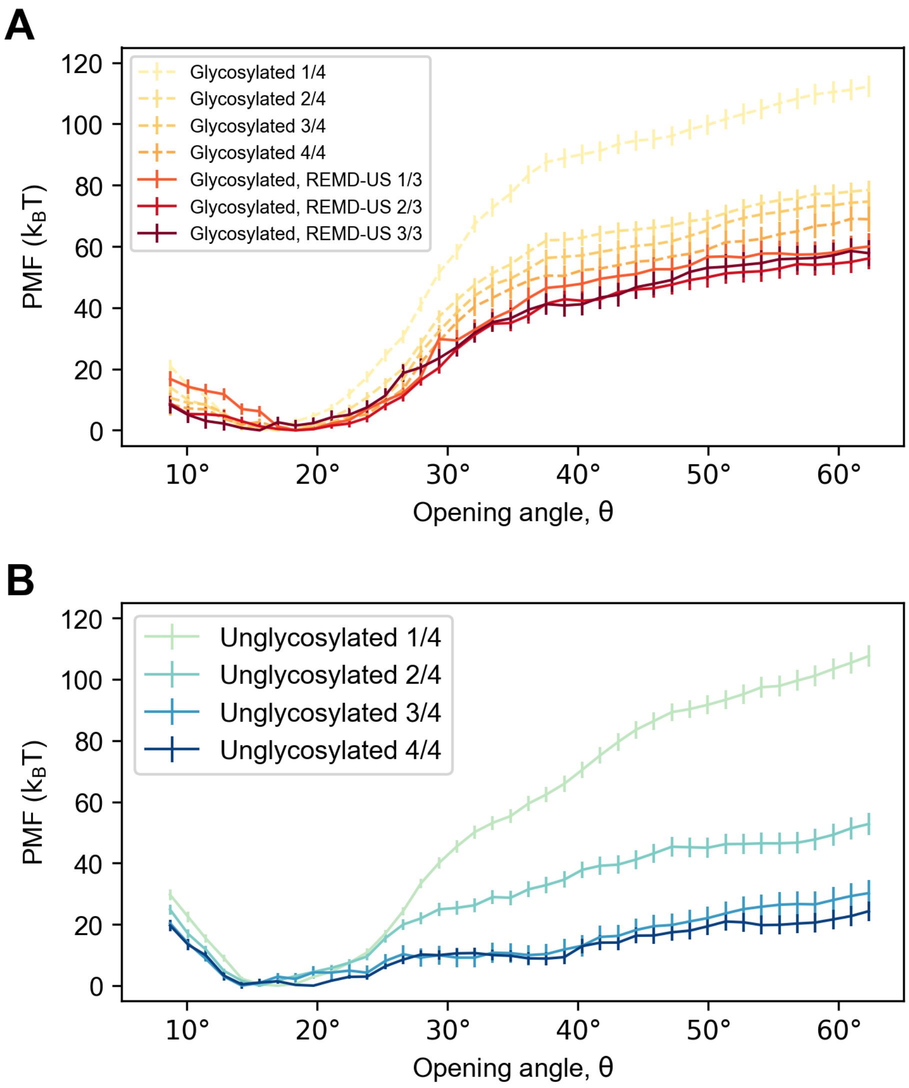

Figure S9: Convergence analysis for the alternative S2-exposure simulations where the RBD was pulled independently of its nearest NTD.
For both panels (A) and (B), different non-overlapping portion of the umbrella sampling trajectories were used to construct PMFs. Fractions denote the portion used (e.g. 2/4 means second of four segments). For the glycosylated system in (A), ordinary umbrella sampling was followed by REMD-US sampling, and the last two thirds of trajectories produce the same PMF. The unglycosylated system in (B) achieved convergence in the second half of ordinary umbrella sampling.

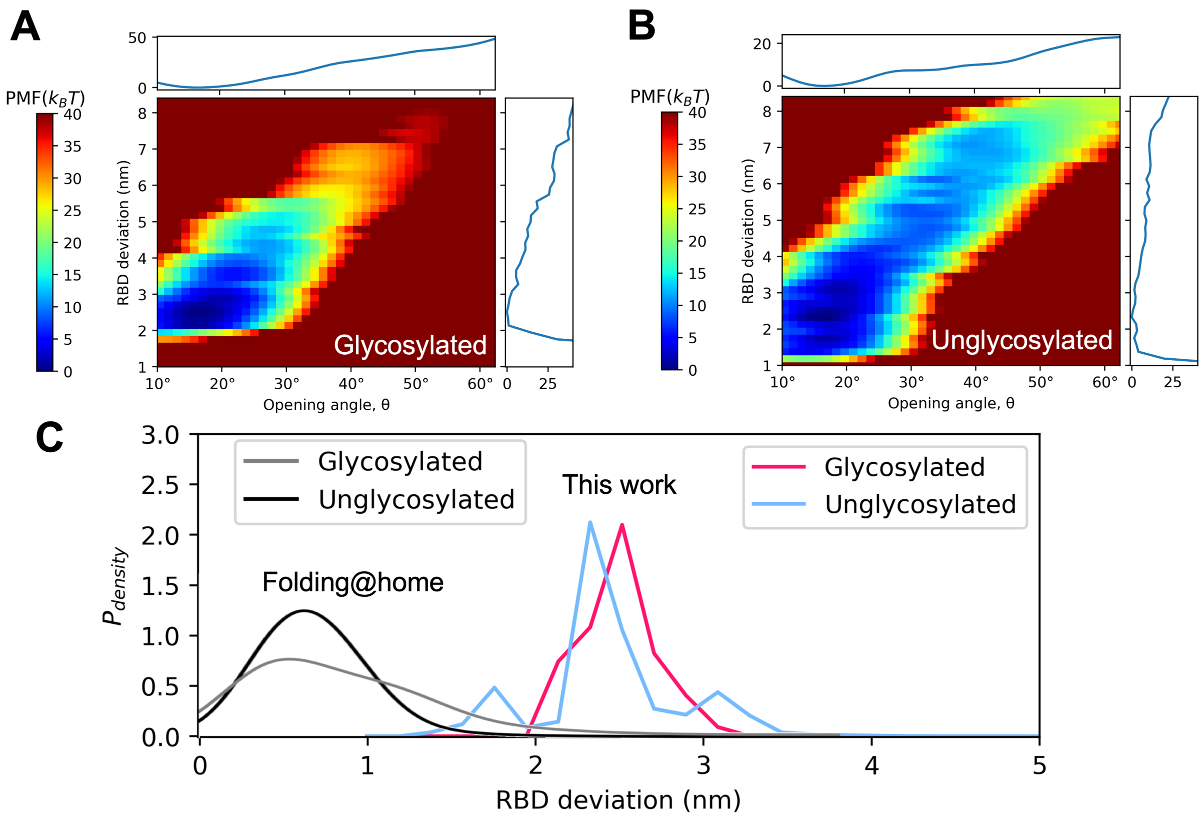

Figure S10: Transformation of the spike opening PMF along RBD deviation order parameters.
In order to compare our S2 exposure PMF against the result obtained from the Folding@home study [3], we transformed our angular coordinate ($\theta$) to the coordinate used in their study, i.e. the distance of the center of mass of the moving RBD from its down state position. For this, first the two-dimensional PMFs of (A) glycosylated and (B) unglycosylated system were constructed along these two order parameters. The RBD deviation of each sampled spike structure was calculated with respect to the position of a virtual down-state RBD which was inferred by assuming that the plane formed by the centers of mass of the two down-state RBDs and the virtual down-state RBD is orthogonal to the spike central helix (i.e. the vector $\vec{PQ}$ in Fig. 1A in the main text). The 2D PMF was constructed using PyMBAR (v4.0.1) module with a kernel size of 0.03 in kernel density estimation (KDE). The 1D projections of this 2D PMF along the two order parameters are shown in units of $k_BT$ along the edges in both panels A and B. (C) Our transformed marginal PMF along the RBD deviation was converted to probability density (dark pink and light blue), and compared to that in Fig. S1 of Zimmerman et. al. [3] (black and grey).

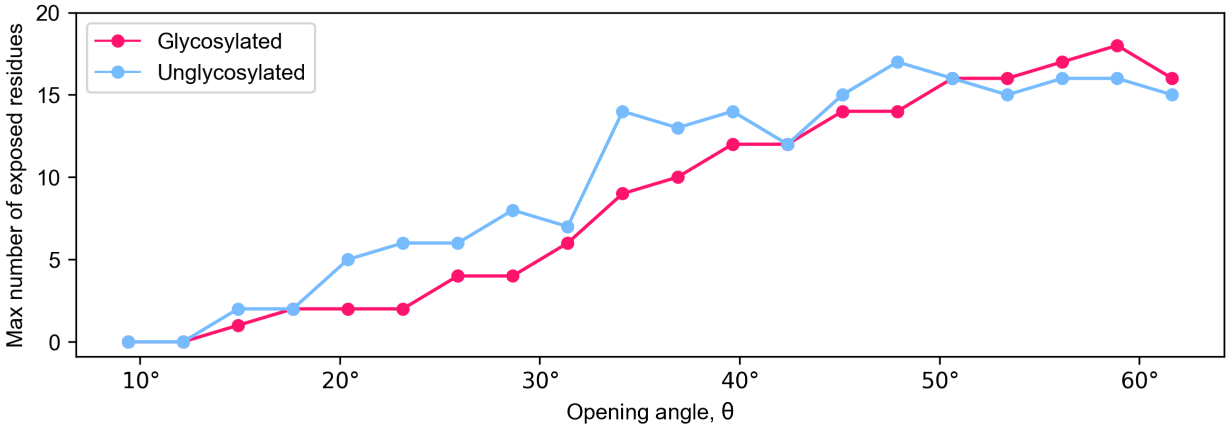

Figure S11: Residue exposure observed in RBD opening simulations.
In each angle bin, the maximum number of residues in the epitope potentially accessible to antibodies is shown. A residue is defined as accessible for antibody binding if its AASA is greater than 1 $\mathring{A}^2$. The binning is the same as that in Fig. 2 of the main text.

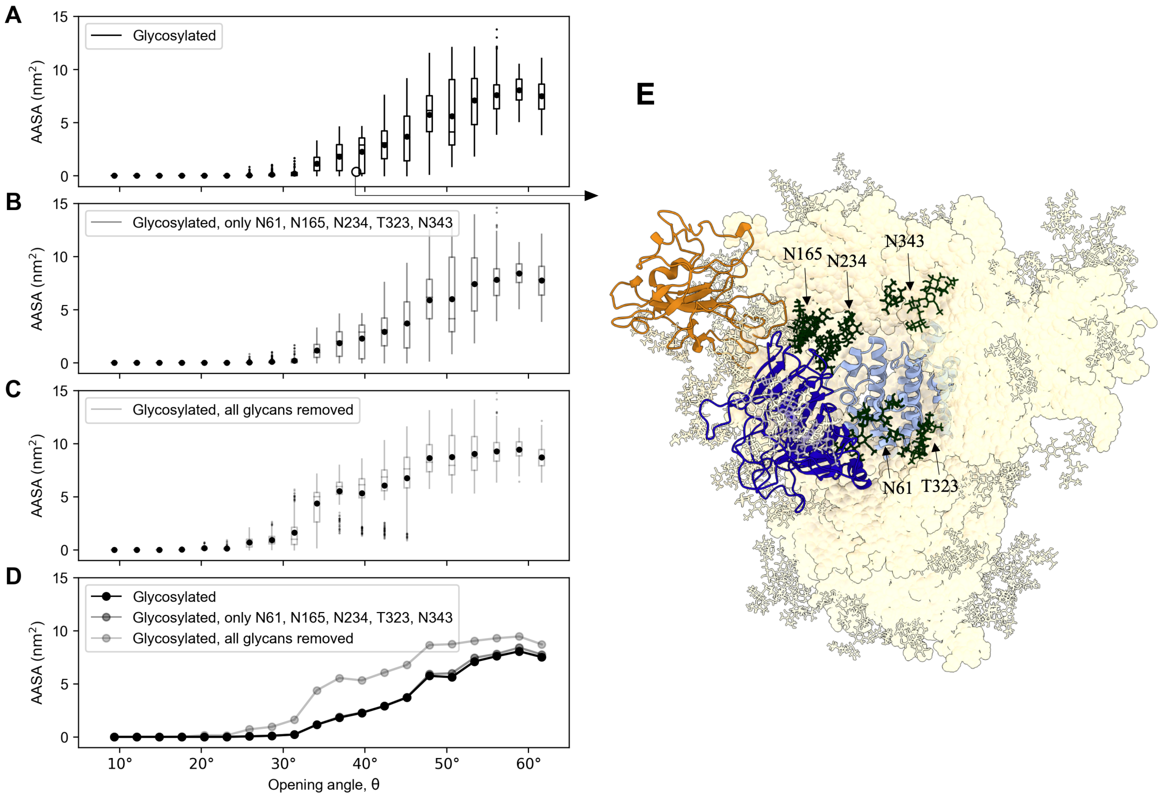

Figure S12: Effect of glycans on the epitope accessibility

The AASA of the S2 region was calculated for the glycosylated spike system accounting for the presence of different sets of glycans. Glycans on Asn-61, Asn-165, Asn-234, Thr-323, and Asn-343 on chain B (i.e the chain that has the NTD closest to the up-state RBD on chain A) contribute the most to protection of the S2 epitope. (A) AASA when all glycans are included. (B) AASA when only glycans on the above selected residues are included. (C) AASA when all glycans are excluded from AASA calculation. (D) The means from panels A-C are plotted together to compare different extents of glycosylation. (E) A snapshot of the fully glycosylated spike at an opening angle of $\theta = 39.0°$ is shown. Its AASA and position along $\theta$ is shown in panel A as an empty circle. The opening RBD is shown in orange. The NTD that moves with the RBD is shown in dark blue. The S2 epitope neighbourhood that the AASA was calculated for is shown in light blue. The selected glycans on Asn-61, Asn-165, Asn-234, Thr-323, and Asn-343 are shown as dark green sticks, and the rest of the glycans are shown as transparent sticks.

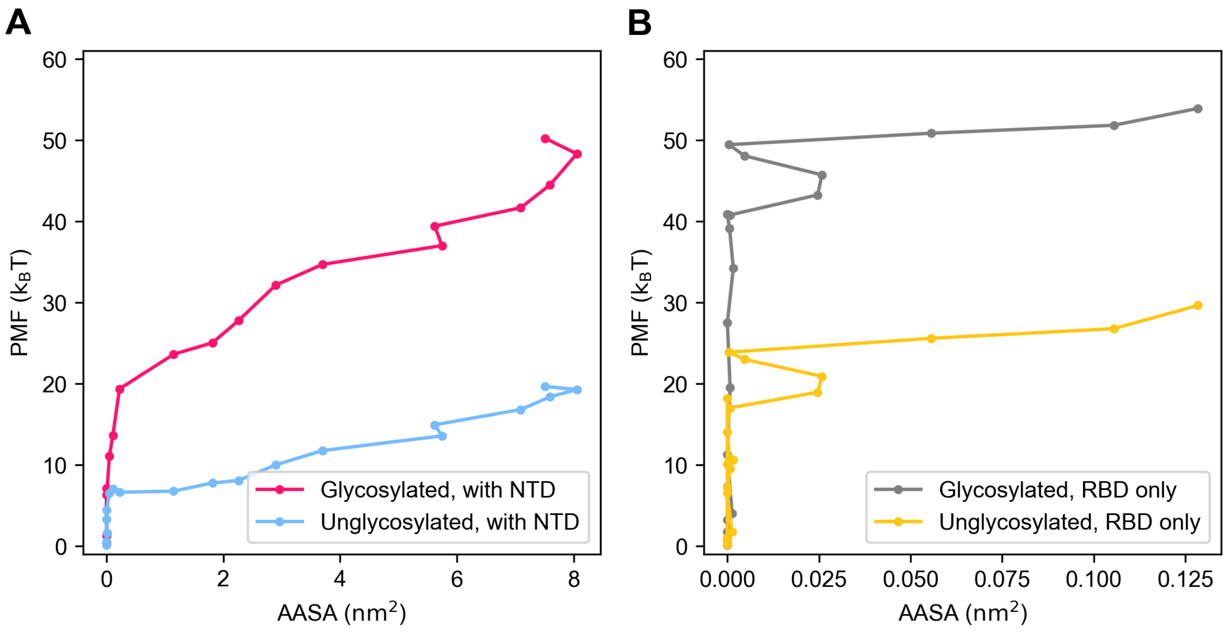

Figure S13: The relationship between AASA and PMF in S2-exposure simulations. This figure uses the same data as Fig. 2 of the main text and Fig. S8 below for panels A and B respectively. The PMF and mean AASA corresponding to each angle bin is shown. PMF values were calculated by linearly interpolating between available data points. Note the very different scales on the x-axes of the two panels.

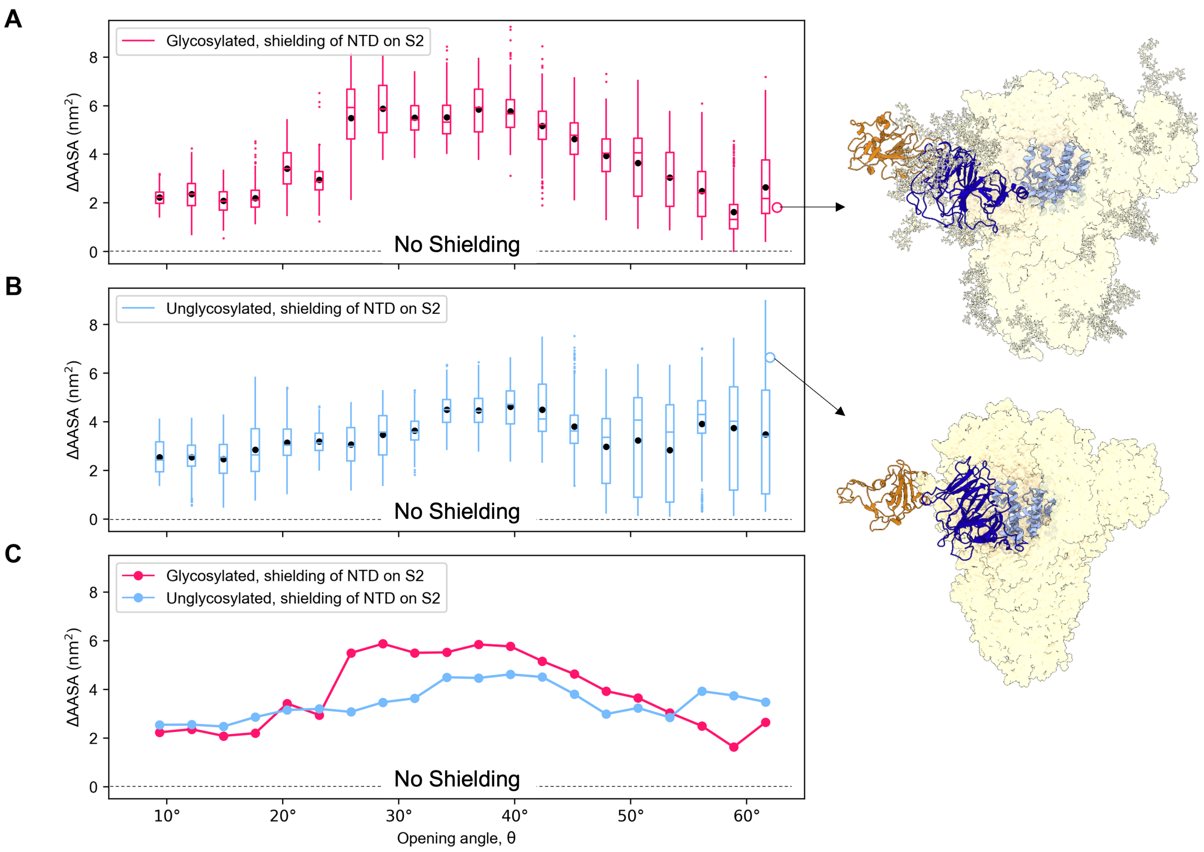

Figure S14: Contribution of the NTD to shielding of the S2 epitope

The difference between AASA of residues 737-765 and 961-1010 in the S2 epitope was calculated with and without the presence of the NTD closest to the opening RBD. The NTD consists of residues 13-305 and their associated glycans. The analysis was performed for both for (A) glycosylated and (B) unglycosylated spike. No shielding by the NTD corresponds to value of $\Delta AASA = 0$ (dotted lines). The upward trend at higher angles in panel B indicates that the NTD provides an increasing degree of shielding at these angles for the unglycosylated system. Representative structures from the two systems are shown on the right with their angular posisions and $\Delta AASA$ values marked on the plots as empty circles. The color rendering is the same as Fig. 1A in the main text. (C) The means from panels A and B are plotted together for comparison of the two systems.

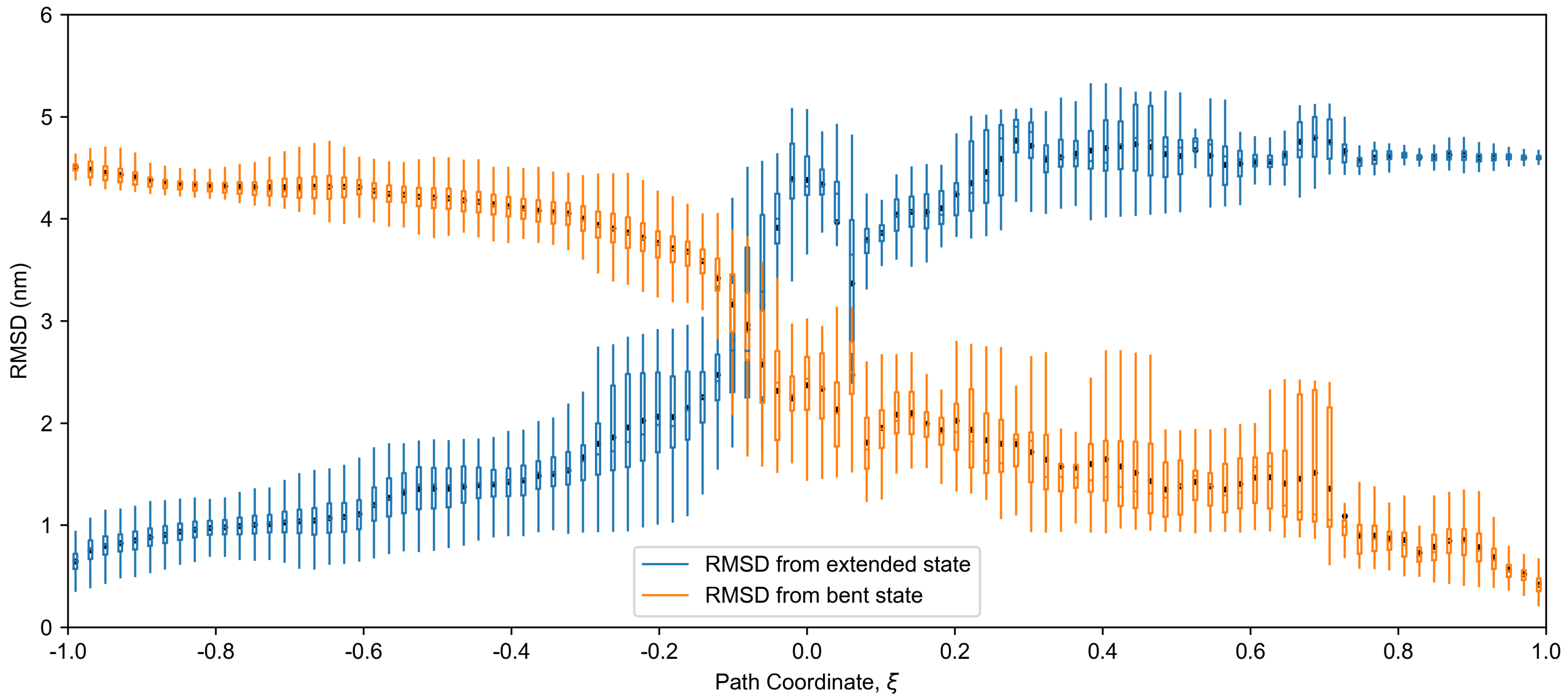

Figure S15: RMSD to end states along the reaction coordinate for helix extension simulations. Frames from the second half of the umbrella sampling trajectories were binned into 100 evenly spaced bins along the coordinate $\xi$, and their RMSD was calculated from the reference structure of each end state. The distribution of the obtained values is shown as a box plot. Means in each bin are shown as black dots, box boundaries correspond to the first and third quartiles, whiskers extend up to the last data-point within 1.5 times the inter-quartile distance, and outliers are not shown.

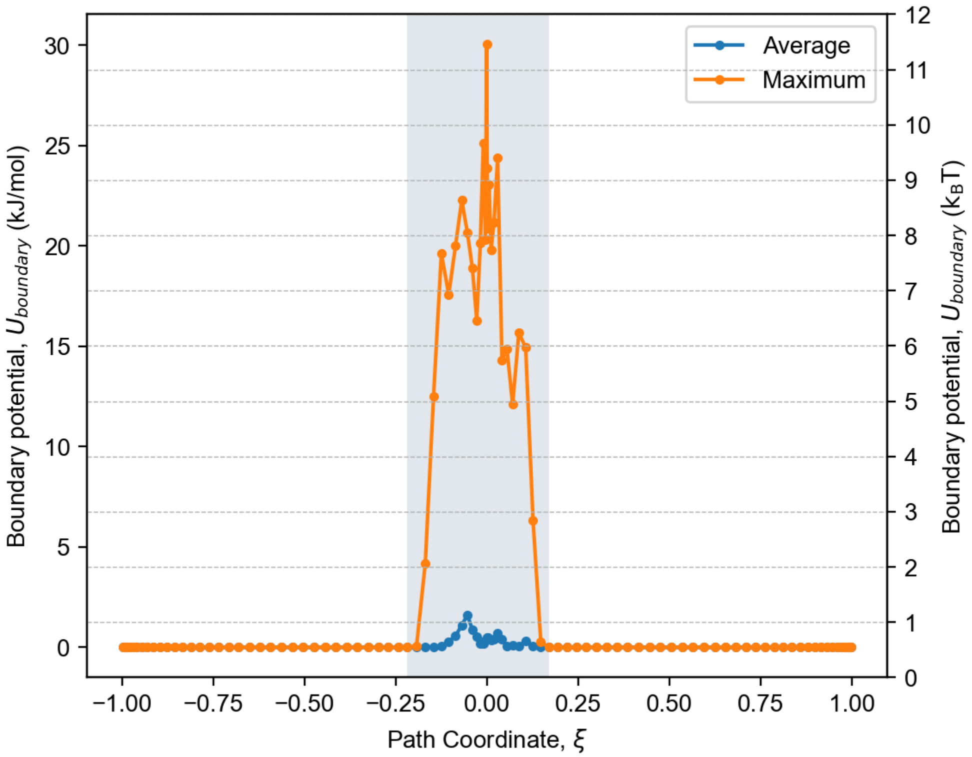

Figure S16: Effect of the boundary potential on helix extension trajectories. The boundary potential experienced in different portions of the phase space during umbrella sampling is shown. Frames were binned by their positions along $\xi$ with umbrella centers set as bin boundaries. The shaded region contains frames with non-zero boundary potential. Also see distribution of frames in $s(X, A)$, $s(X, B)$ space in Fig. 4 of the main text.
